# Short-interval reburns in the boreal forest alter soil bacterial communities, reflecting increased pH and poor conifer seedling establishment

**DOI:** 10.1101/2021.03.31.437944

**Authors:** Jamie Woolet, Ellen Whitman, Marc-André Parisien, Dan K. Thompson, Mike D. Flannigan, Thea Whitman

## Abstract

Increasing burn rates (percentage area burned annually) in some biomes are leading to fires burning in close succession, triggering rapid vegetation change as well as altering soil properties. Despite the importance of soil microbes for nutrient cycling and as plant symbionts, the effects of increased fire frequency on belowground microbial communities remain largely unknown. We present a study of the effects of short interval reburns (defined here as <20 years between fires) on soil bacterial communities in the boreal forest of northwestern Canada, using a paired site design that spans wetlands and uplands, with 50 sites total. We asked whether short interval reburns significantly alter soil bacterial community composition and richness, and which bacterial taxa are associated with greater or lower fire frequency. We found that, while short interval reburns had no significant effect on bacterial richness, there were significant changes in overall community composition. We did not find correlations between understory vegetation community dissimilarities and bacterial community dissimilarities, suggesting the primary drivers of changes induced by short interval reburns may differ between plants and microbes. We identified an abundant *Blastococcus sp*. that was consistently enriched in short interval reburns, in both wetlands and uplands, indicating its role as a strongly “pyrophilous” bacterium. We also identified an abundant *Callaberonia sordidicola* taxon as being consistently depleted in short interval reburns. This endophytic diazotrophic organism is a robust colonizer of pine and spruce seedlings and has the ability to increase seedling growth, due in part to large contributions of fixed nitrogen. Its depletion in short-interval reburn sites raises questions about whether this is contributing to – or merely reflects – poor conifer seedling recolonization post-fire at short-interval reburns.

## Introduction

The boreal zone is one the world’s largest biomes, spanning 1.89 billion ha across the northern hemisphere (Brandt *et al.,* 2013). This zone consists of forests of cold-tolerant tree species, lakes, rivers, wetlands, and naturally treeless areas, such as shrublands and grasslands (Brandt, 2009). The Canadian boreal forest represents 28% - about 552 million ha - of the world’s boreal zone (Brandt *et al.* 2013) and provides habitat for thousands of species, supplies numerous ecosystem services including timber, timber products, and water filtration, is home to 12% of Canada’s population, and offers many other economic and cultural resources (Bogdanski, 2008). Furthermore, this system stores about 10 - 30% of the global terrestrial carbon stocks, mostly belowground in peatlands and soils (Bradshaw and Warkentin, 2015; Kasischke, 2000), which may be threatened by changing fire regimes (Ribeiro-Kumara *et al.*, 2020, Walker *et al.*, 2019).

Fire is a common and widespread disturbance throughout much of the western Canadian boreal zone, where the average fire-free interval has been observed to range anywhere between 30 – 100s of years (Larsen 1997; Stocks 2002). Fire is a critical event for maintaining healthy boreal ecosystems by shaping vegetation composition, soil chemical properties, and animal communities (Rowe and Scotter, 1973). Over the past 50 years, there has been a shift in the forest fire regime for many areas of the North American boreal forest, including lengthened burn season, increased lightening ignitions, and increased area burned (Hanes *et al*., 2019; Wotton and Flannigan, 1993; Jain *et al*., 2017; Veraverbeke *et al.,* 2017). These shifting disturbance regimes can have adverse effects on ecosystems and may degrade forest resilience to fire (Johnstone *et al*., 2016). Forest resilience, broadly, is the ability for a forest to return to pre-disturbance conditions, often determined by ecological memory of past states (*e.g.*, via seed banks) (Johnstone *et al*., 2016) and the regeneration of plant communities (Gill *et al*., 2017). Under normal fire regimes, forests often have self-regulatory processes that limit disturbance frequency (Peterson, 2002). Under drought conditions and as the forest ages, however, these self-regulatory processes can weaken, allowing for increased fire frequency (Parks *et al.,* 2018). In the relatively uncommon case when young forests (<20 years) reburn, increased fire frequency (short interval reburns) in boreal forests can alter vegetation composition (Whitman *et al*., 2019b), change above-ground plant production (Johnstone and Chapin, 2006), and potentially induce forest-type conversions (*i.e.*, *Picea* spp.-dominated to *Populus tremuloides*- or *Pinus banksiana-*dominated) (Johnstone and Chapin, 2006; Gill *et al,* 2017). Short interval reburns also reduce soil organic horizon thickness (Johnstone and Chapin, 2006; Hoy *et al.,* 2016), change soil chemical properties by depleting total C and N (Pellegrini and Jackson, 2020; Pellegrini *et al.*, 2020), and can potentially decrease microbial decomposition rates (Köster *et* al., 2015; Pellegrini *et al.*, 2020). Furthermore, novel wildfire regimes have been found to have long-term effects on biogeochemical soil processes, decreasing mineral soil organic carbon (SOC), soil extracellular enzyme activity, and soil microbial respiration (Dove *et al.,* 2020), which may interact with vegetation responses to fire (Knelman *et al*., 2015). However, it is less clear how short interval reburns may affect soil microbial communities.

Soil microbial communities provide numerous critical ecosystem functions, including cycling carbon and nitrogen, supporting plant growth and diversity, preventing erosion, and maintaining soil structure via biofilms and fungal hyphae (Van Der Heijden *et al.,* 2007; Saleem *et al.,* 2019). Fire can affect the soil microbial communities directly, via heating and oxidation of the soil environment, and indirectly, by increased exposure to climatic variation and from changes to the physicochemical environment (Hart *et al.*, 2005). Immediately post-fire, microbial biomass may decrease due to direct killing of microbes or the loss of nutrient resources (Dooley and Treseder, 2011; Holden and Treseder, 2013; Pressler *et al*., 2019). Microbial community structures may take decades to recover to previous states, often requiring plant community reestablishment first (Dooley and Treseder, 2011; Ferrenberg *et al.,* 2013). The extent to which microbial community composition is affected by fire is influenced by fire severity and changes to vegetation, moisture, pH, and soil carbon after fires (Whitman *et al*., 2019a; Hart *et al*., 2005; Holden and Treseder, 2013; Sáenz de Miera *et al*., 2020).

Microbial resistance and resilience may inform our understanding of forest resilience following fire. Microbes have innate traits that differ from those of some of their larger-organism counterparts (high abundances, widespread dispersal potential, comparatively rapid growth potential, and comparatively rapid evolutionary adaptations (Shade, 2012)).The stability of their populations over time can be influenced by an individual’s stress tolerance and phenotypic plasticity, a population’s growth rate and adaptability, and a community’s richness, evenness, and microbial interactions (Shade, 2012). Microbial communities may show resistance by recovering as a compositionally different community, yet remaining functionally similar, in that the ecosystem process rates of interest remain unchanged (Allison and Martiny, 2008). Short interval reburns have the potential to change soil properties and vegetation community composition, both of which are factors that shape microbial communities (Chandra *et al.,* 2016; Woolet and Whitman, 2020; Van Der Heijden *et al.*, 2007; Bardgett and van der Putten, 2014). However, there are few studies that examine how microbial communities respond to these potentially transformative short interval reburns. Because microbes play a critical role in ecosystem functioning and structure, it is important to understand how they respond to fire regime changes.

Using a paired sample study design, Whitman *et al*. (2019b) showed that changes in fire-return interval caused changes to conifer and broadleaf recruitment, soil organic horizon depth, and herbaceous vegetation cover. Here, we set out to investigate soil bacterial community responses to paired short interval (SI) vs. long interval (LI) reburns at the same sites (Whitman *et al.* 2019b). Specifically, we asked 1) Do microbial communities have a different response to short interval reburns compared to normal fire intervals?, 2) Do short interval reburns reduce bacterial community richness?, and 3) Which bacterial taxa respond positively to increased fire frequency? We hypothesized that:

1. There would be a significant effect of SI vs. LI reburns on bacterial community composition, where the (dis)similarity of bacterial communities between paired SI and LI sites would be dependent on:

a. Time since last fire (TSLF): paired sites that had a longer time to recover at the sampling date would be more similar than sites that had less time to recover
b. Vegetation transition: paired sites that show a change in leading tree species would be less similar than pairs that did not see a change in leading tree species due to changes in rhizosphere interactions and differences in litter inputs
c. Drainage class: paired wetland sites would be more similar than paired upland sites due to wetland sites generally having lower severity fires and thus retaining more of the organic horizon
d. Difference in fire-free interval (FFI): paired sites that had greater differences in fire-free interval would be less similar due to larger pre-fire differences in successional trajectory
2. Bacterial richness would not be consistently different between paired SI and LI sites, but would be predicted by pair-specific characteristics (TSLF, difference in FFI)
3. Bacteria that have higher relative abundance in SI sites than LI sites would include taxa that have previously been identified as being responsive to fires and associated with increasing burn severity in this region, such as certain taxa within the genera *Blastococcus*, *Arthrobacter*, and *Massilia* (Whitman *et al*., 2019a).

## Methods

### Study region description

Our study area is located in the Canadian boreal forest in the Northwest Territories and Alberta around Tu Nedhé (Great Slave Lake) (Figure 1).

**Figure 1.**
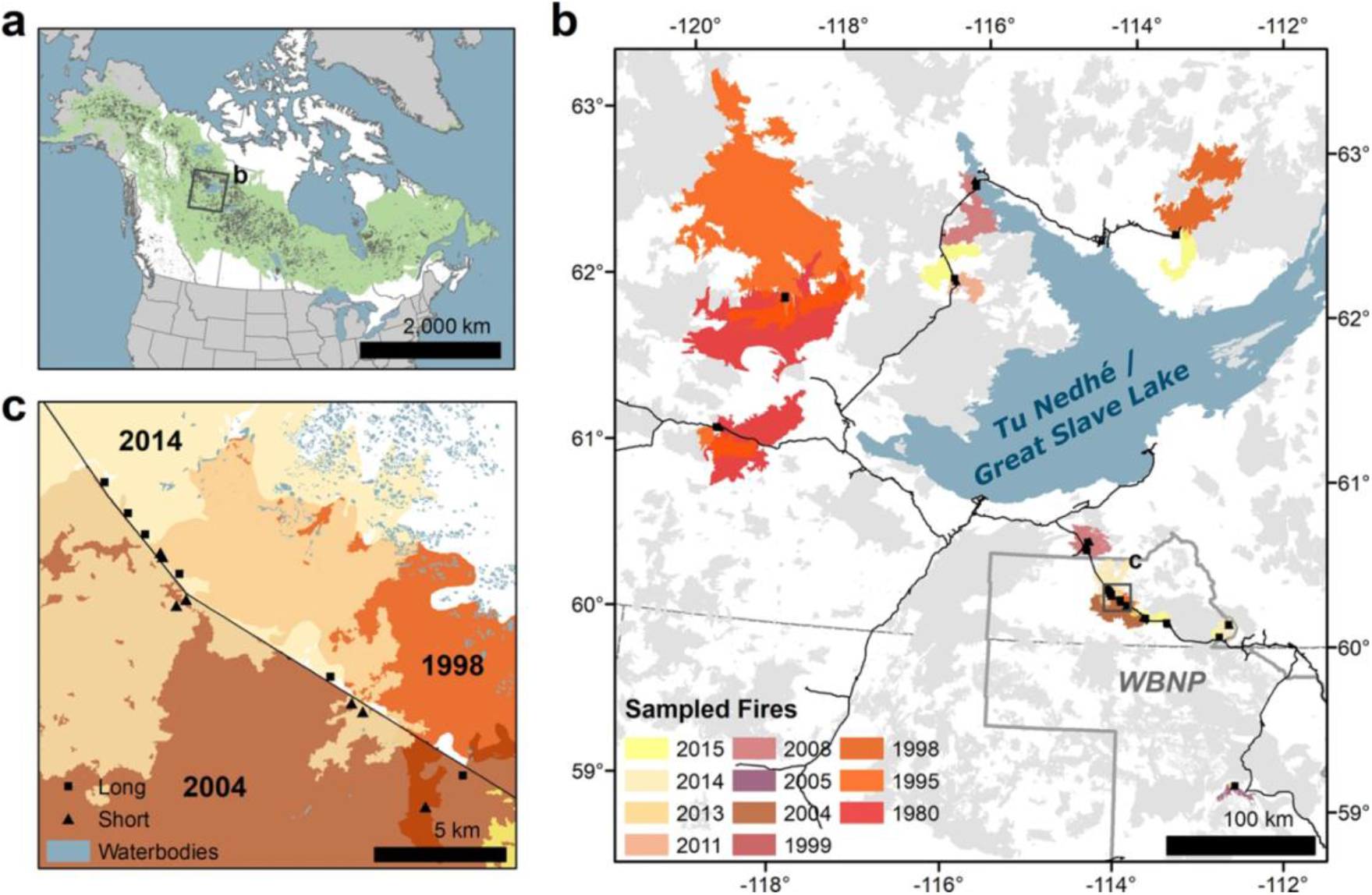
Location and fire history of the study area (Figure adapted from Whitman *et al*., 2019b). (**a**) Recent (1984–2016) fire perimeters (dark grey) within the boreal forest of North America (green). (**b**) Fire history within the study region around Tu Nedhé (Great Slave Lake). Recent wildfires (1980–2015) are coloured grey, while sampled wildfires are coloured by year of occurrence. Shapes represent sampling locations. The boundary of Wood Buffalo National Park (WBNP) is outlined in grey and major roads are shown in black. (**c**) Detail showing an example of the sampling design of paired sites with short (triangles) and long (squares) interval sites. E.g., consider the triangle (SI) and square (LI) in SW corner – both were burned in the 2004 wildfire, but only the SI site was also burned in the 1998 wildfire.

Typically, the climate consists of short, warm summers, and long, cold winters. Fire frequencies in this area range from hundreds of years to 30 years between stand-replacing fires (Larsen and MacDonald, 1998; Stocks *et al.* 2002). The landscape consists of open wetlands, forested peatlands, and forested uplands, with black spruce (*Picea mariana* (Mill.) Britton, Sterns & Poggenb.), white spruce (*Picea glauca* (Moench) Voss), jack pine (*Pinus banksiana* Lamb.), and trembling aspen (*Populus tremuloides* Michx.) as the dominant tree species. The soils in this region are mostly classified as Mesisols, Gleysols, and Luvisols. The sample sites span a wide range of soil properties, with pH values ranging from 5.13–8.11 in the mineral horizons and 5.29–8.35 in the organic horizons, total C ranging from 0.4% (mineral horizon) to 44.6% (organic horizon), and a wide range of textures (Table 1). Detailed information on the study area and field sampling methods can be found in Whitman *et al.* (2019b).

**Table 1.**
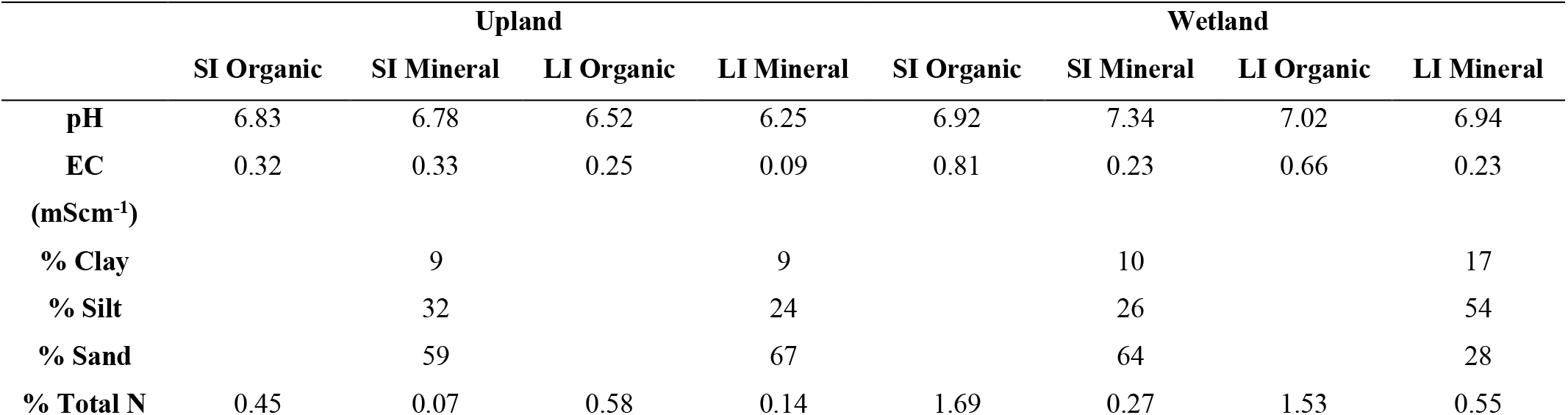
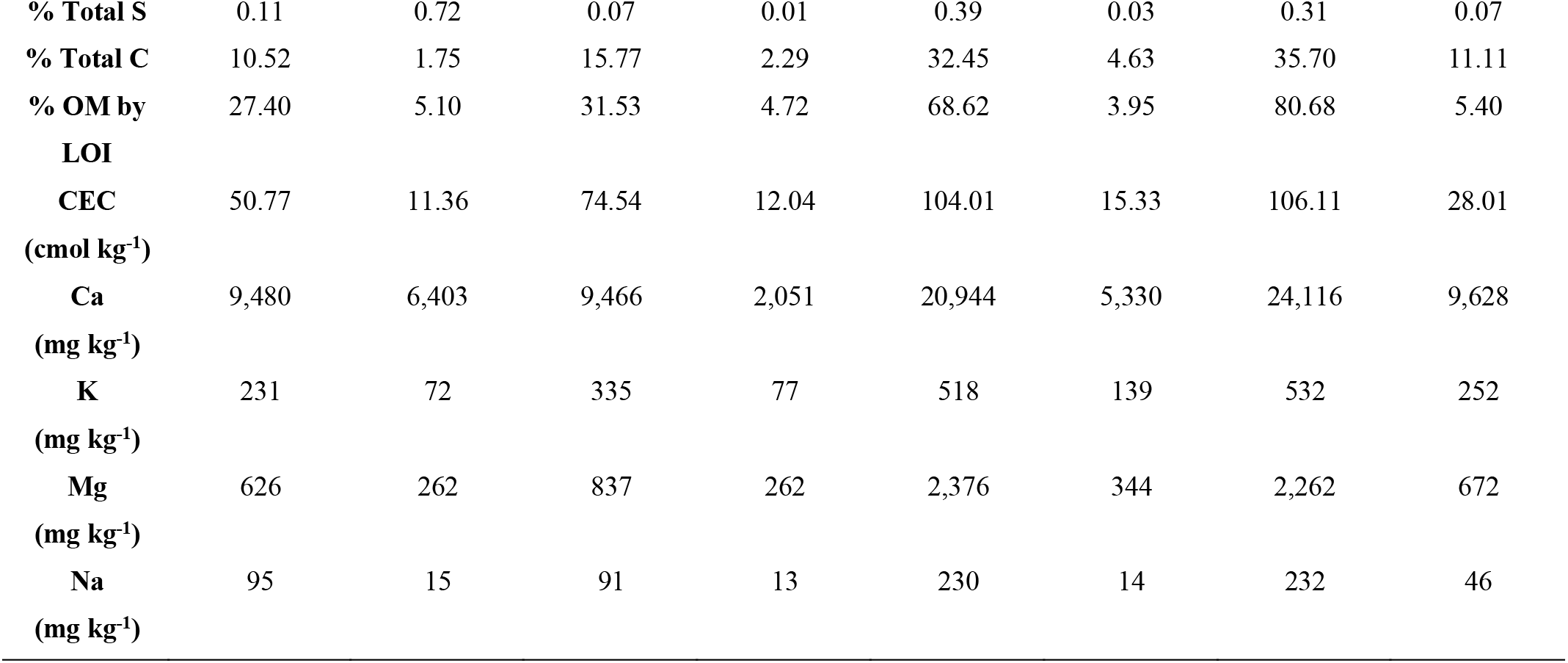
Soil properties across drainage class, short and long fire-free intervals, and organic and mineral soil horizons. Means reported (N = 3). SI=Short-interval, LI=Long-interval.

### Experimental design and site assessment

In 2016, 50 sites (25 pairs) were sampled; eight of these pairs were wetlands and 17 were uplands. Paired sites shared the same drainage class, fire history, and their most recent fire, which burned between 1995 and 2015, but had different previous fires, which allowed us to characterize them as short fire-free interval fires (SI; fire-free interval was 4-17 years) or long fire-free interval fires (LI; fire-free interval was 30-112 years). Fire history maps and fire-scarred trees were used to date recent and previous fires (Whitman *et al*., 2019b). Time since last fire at sampling varied across the dataset, ranging from 1 – 21 years.

Sites were selected > 100 m from roads. At each site, a 35 m transect oriented north-south was used to collect vegetation data and soil samples. Vegetation survey methods are described in detail in Whitman et al. (2019b). Briefly, along the transect, overstory structure was assessed by sampling live and dead mature trees (greater than 1.3 m in height and greater than 3 cm diameter at breast height (DBH)) stem density, basal area, and species composition. Understory structure was assessed by collecting vegetation and small shrub abundance, tall shrub counts, the density, species, and status (live or dead) of seedlings and saplings, surface fuel, and coarse woody debris. At 0, 17.5, and 35 m along the transect, soil cores (5.5 cm diameter, 13.5 cm depth) were sampled by gently extruding and separating the core into organic (O) horizons (where present, up to 13.5 cm) and mineral (M) horizons (where present, up to 5 cm). Wetland mineral soils underlying thin organic horizons were only present within the top 13.5 cm at three sites. The three samples from the transect were pooled by horizon at each site and mixed gently by gloved hand in a bag. From these site-level samples, sub-samples were collected for microbial community analysis and stored in LifeGuard Soil Preservation solution (Qiagen, Germantown, MD) in a 5 mL tube. Tubes were kept as cold as possible while in the field, then stored frozen. The remaining soil samples were air-dried and analyzed for soil properties, as described in detail in Whitman *et al*. (2019b) (Table 1).

### DNA extraction, amplification, and sequencing

DNA extractions were performed for each sample, with a blank extraction every 24 samples, using a DNEasy PowerLyzer PowerSoil DNA extraction kit (QIAGEN, Germantown, MD) following manufacturer’s instructions. LifeGuard Soil Preservation solution was removed from samples by thawing the sample on ice and centrifuging for 2 minutes at 10,000 x g. Preservation solution was gently pipetted off to remove as much as possible, and the sample was centrifuged a second time for 30 seconds at 10,000 x g to remove any remaining preservation solution. The sample was re-mixed with a spatula before weighing for extraction. Extracted DNA was amplified in triplicate PCR reactions, targeting the 16S rRNA gene v4 region (henceforth, “16S”). Reactions were performed in 96-well plates with each PCR mixture containing 12.5 μL Q5 Hot-Start High-Fidelity 2X Master Mix (New England Biolabs, Ipswich, MA), 1.25 μL 10 μM 515F primer (AATGATACGGCGACCACCGAGATCTACAC-barcode-TATGGTAATT GTGTGYCAGCMGCCGCGGTAA), 1.25 μL μM 806R primer (CAAGCAGAAGACGGCATACGAGAT-barcode-AGTCAGCCAGCCGGACTACNVGGGTWTCTAAT) (Walters *et al.* 2015; Kozich *et al.* 2013), 1.25 μL 20 mg mL^−1^ BSA, 7.75 μL nuclease-free water, and 1 μL DNA template. Positive control (bacterial isolate DNA) and negative control (nuclease-free water) reactions were included on each plate. PCR mixtures were amplified on an Eppendorf Mastercycler nexus gradient thermocycler (Eppendorf, Hamburg, Germany) at the following conditions: 98°C for 2 minutes, (98°C for 10 seconds, 58°C for 15 seconds, 72°C for 10 seconds) x 30 cycles, 72°C for 2 minutes, hold at 4°C. PCR amplification success was verified via gel electrophoresis on a 1% agarose gel. The amplicon triplicates for samples and extraction blanks were pooled and normalized using a SequalPrep Normalization Plate (96) Kit (ThermoFisher Scientific, Waltham, MA), following manufacturer’s instructions. Normalized samples were pooled and library cleanup was performed using a Wizard SV Gel and PCR Clean-Up System A9282 (Promega, Madison, WI). The purified library was submitted to the UW Madison Biotechnology Center (UW-Madison, WI) for 2×250 paired end (PE) Illumina MiSeq sequencing.

### Sequence data processing and taxonomic assignments

The University of Wisconsin – Madison Biotechnology Center performed demultiplexing on sequences. The total read count was 5,141,495 sequences; the minimum (non-blank) read count was 18,297; the maximum read count was 52,378; and the mean read count (not including blanks) was 30,826; sample blanks averaged 2,437 reads and did not produce any visible bands in electrophoretic gels. Forward and reverse reads were imported into a Jupyter Notebook where QIIME2 (QIIME2, v 2019.10) and dada2 (Callahan *et al*., 2016) were used to filter, learn error rates, denoise, and remove chimeras, to generate operational taxonomic units (OTUs) – specifically, amplicon sequence variants. After quality control steps, a total of 3,593,901 reads were retained. Taxonomy was assigned using the QIIME2 scikit-learn feature classifier trained on the 515f-806r region of the 99% ID OTUs (Bokulich *et al.*, 2018) from the Silva138 database (Quast *et al*., 2013). All sequences are deposited in the NCBI SRA under accession numbers [to be submitted upon paper acceptance].

### Statistical analyses

Analyses and plotting were done with R (R Core Team, 2019) in Jupyter notebooks and RStudio, using packages *phyloseq* (McMurdie and Holmes, 2013), *dplyr* (Wickham et al., 2019), and *ggplot2* (Wickham, 2016).

To test for the effect of reburn interval on bacterial community composition, we calculated the Bray-Curtis dissimilarities for all sites from relative abundances, using the *vegan* package in R (Oksanen et al., 2018, Bray and Curtis, 1957), and used permutational multivariate analysis of variance (PERMANOVA) to determine whether SI *vs*. LI was a significant predictor of community composition after controlling for paired sites. We plotted the dissimilarities using NMDS and used the *envfit* function in *vegan* to map site characteristics onto the ordination. To determine what factors to control for in subsequent data analyses, we used PERMANOVA to test whether drainage class, soil horizon, and pH were also significant predictors. We then calculated Bray-Curtis dissimilarities for each site-horizon pair. After controlling for soil horizon and drainage class (except for when testing drainage class itself), we used ANOVA to test whether the dissimilarities between paired sites were affected by 1) time since last fire, 2) vegetation transition (*i.e.*, whether there was a change in leading species status, as inferred from different leading species in the paired sites), 3) drainage class (wetland or upland), and 4) difference in FFI between paired sites.

Because vegetation communities were affected by FFI in Whitman *et al.* (2019b), we also examined the relationship between vegetation-related parameters and bacterial community dissimilarities in LI *vs*. SI sites. These parameters included the effects of absolute difference in density of small understory tree stems between paired sites (including all stems, broadleaf stems, and conifer stems), difference in percent understory vegetation cover, and Bray-Curtis dissimilarities of understory vegetation community. Additionally, in some figures, we indicate sites where fires were likely to have been low-severity surface fires. This rough designation was based on whether sites had live trees with large mean DBH (>5 cm), excluding sites where mean height was <= 4 m, and TSLF >= 5 years, as trees could grow to this height/DBH in that time.

In order to determine whether bacterial richness was affected by short interval reburns, we estimated the richness of bacterial communities by using the R package *breakaway* (Willis and Bunge, 2015), using a weighted linear regression model (Rocchetti *et al*., 2011). To test for differences in richness between paired SI and LI samples in the organic and mineral horizons, we used paired *t*-tests. We then calculated the percent difference in richness estimates between paired SI and LI sites and used an ANOVA to test for the effect of TSLF, vegetation transition, drainage class, difference in FFI, and absolute difference in total number of stems, controlling for soil horizon and drainage class (except for when testing drainage class itself). We also tested whether including bacterial richness improved predictions of post-fire seedling recruitment by using ANOVA, with live overstory stems, site moisture, TSLF, and interval as predictor variables for the first model (as in Whitman *et al*. (2019b)), and live overstory stems, site moisture, TSLF, interval, and bacterial richness in the second model.

In order to determine which specific taxa were associated with LI *vs*. SI sites, we estimated the differential abundances of taxa in LI *vs.* SI sites using the R package *corncob* (Martin *et al.,* 2020) with the Wald testing procedure and a false discovery rate cutoff of 0.05, controlling for sample pair and soil horizon, and analyzing wetlands and uplands separately. Using the differential abundance estimates, we calculated the estimated log_2_-fold change in the relative abundances of the significant taxa in the LI *vs.* SI sites. To identify responding OTUs that were also identified in Whitman *et al.* (2019a), where we studied bacterial and fungal responses to fire severity in the Canadian boreal forest at sites from the same region, we used BLAST (Camacho *et al*., 2009). Every interval-responsive OTU from this study matched an OTU from the previous study at 99.6-100% ID and all were classified according to their log_2_-fold change responses in burned *vs.* unburned sites from the previous study: enriched in burned sites (positive), depleted in burned sites (negative), or no significant change (neutral).

## Results

### Effects of short interval reburns on bacterial community composition

Short interval reburns, relative to long interval reburns, significantly affected bacterial community composition (PERMANOVA, p = 0.002, Figure 2, Supplemental Table 1), after controlling for pair ID. Drainage class (p = 0.001, Figure 2, Supplemental Table 2) and soil horizon (p = 0.001, Figure 2, Supplemental Table 2) were also significant drivers of bacterial community dissimilarities in the dataset, whereas pH was not (p = 0.28, Supplemental Table 2). For the remaining analyses, mineral and organic soil horizons and wetlands and uplands were analyzed separately or were controlled for by including these variables in regressions.

**Figure 2.**
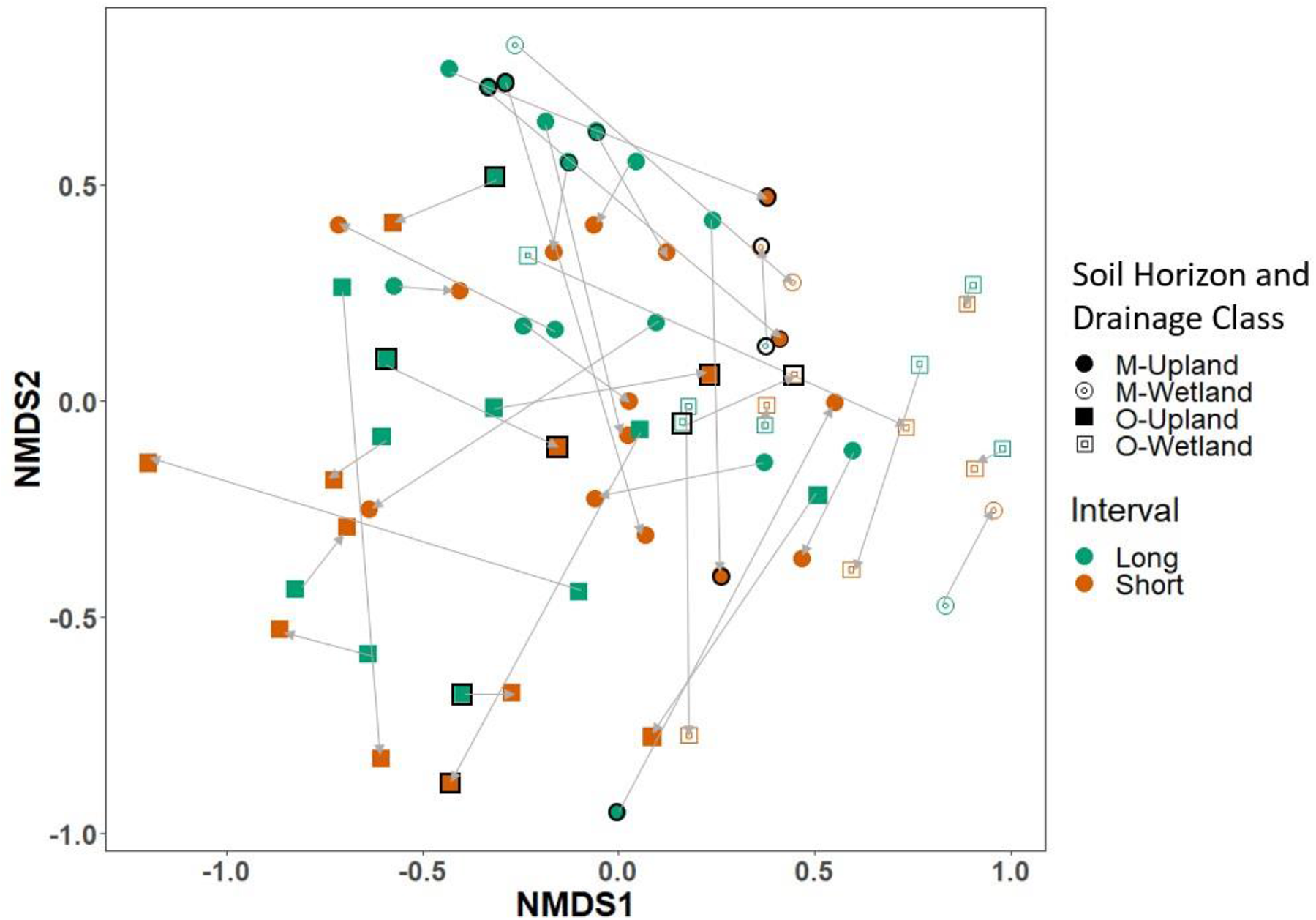
Non-metric multidimensional scaling plot of Bray-Curtis dissimilarities of the bacterial community composition (k = 4, stress = 0.08, first two axes shown). Drainage class is identified by fill (wetland = open symbols, upland = filled symbols), and soil horizon is identified by shape (mineral (M) = circle, organic (O) = square). Reburn interval status is indicated by color (long = green, short = orange). Light grey arrows connect site pairs, pointing from long to short fire-free intervals. Symbols outlined in black indicate samples where fire was characterized as a low-severity surface fire.

The second and fourth NMDS axes were most clearly associated with differences in LI and SI sites. SI sites were associated with a shift away from conifer-dominated to broadleaf-dominated and were also associated with higher soil pH (Figure 3).

**Figure 3.**
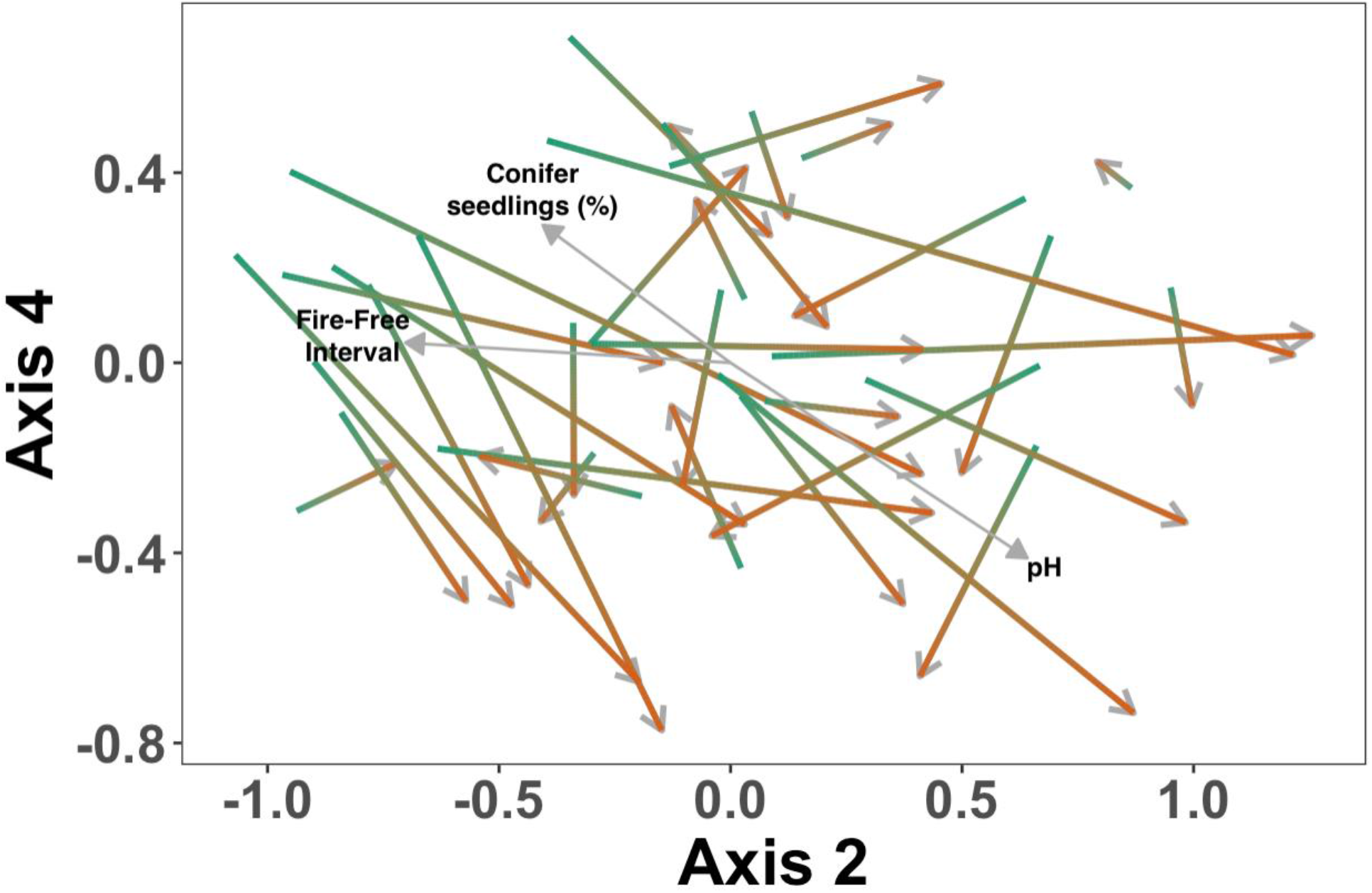
Second and fourth axes of non-metric multidimensional scaling of Bray-Curtis dissimilarities for bacterial community composition (k = 4, stress = 0.08). Paired short and long interval sites are joined by thicker shaded arrows (green blunt end indicating LI site, orange pointed end indicating SI site). Light grey arrows indicate vectors for selected significant (p<0.05) site characteristics – pH, % conifer seedlings, and fire-free interval.

None of our factors of interest had significant effects on bacterial community dissimilarities between paired SI and LI sites. There was not a significant effect of TSLF on dissimilarities between bacterial communities from SI and LI sites (p = 0.19; Supplemental Figure 1 and Supplemental Table 3). Whether there was a vegetation transition between paired SI and LI sites was also not a significant factor driving bacterial community dissimilarities (p = 0.71; Supplemental Figures 1 and 2, Supplemental Table 3). Bacterial community dissimilarities between paired SI and LI sites were not different between wetlands and uplands (p = 0.13; Supplemental Figures 1 and 2, Supplemental Table 3). Lastly, bacterial community dissimilarities between paired SI and LI sites were not clearly affected by differences in FFI (p = 0.12; Supplemental Figure 2 and Supplemental Table 3).

Paired sites where the LI site had more total understory stems and a higher proportion of conifer stems than the SI site had more dissimilar bacterial communities (Figure 4, Table 2). Dissimilarities between bacterial communities in paired SI and LI sites were not affected by difference in total understory vegetation cover (p = 0.91; Supplemental Figure 3, Supplemental Table 4) or vegetation community dissimilarities (R^2^ = −0.01, p-value = 0.96, Supplemental Figure 4; Supplemental Table 4).

**Figure 4.**
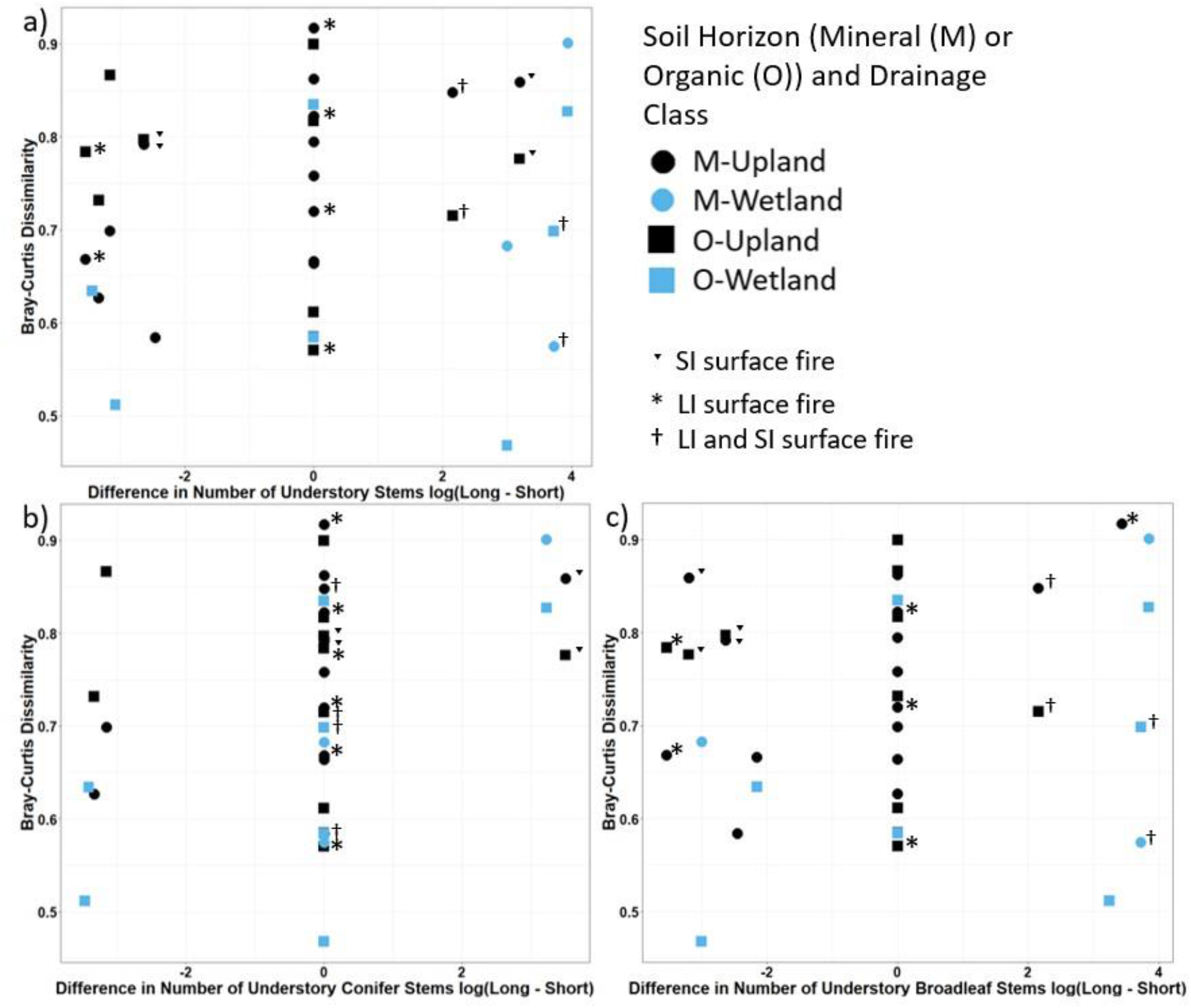
Bray-Curtis dissimilarities of bacterial community composition between paired short interval (SI) and long interval (LI) reburn sites *vs*. the log difference (log(LI - SI) of a) all understory stems per hectare, b) understory conifer stems per hectare, and c) understory broadleaf stems per hectare. Drainage class is identified by color (wetland = light, upland = dark), and soil horizon is identified by shape (mineral (M) = circle, organic (O) = square). Note log scale on x-axis. Values at zero had no stems recorded at time of sampling, and therefore had no differences in stem count. Pairs where the short interval fire was characterized as a low-severity surface fire are indicated by a ˑ symbol, pairs where the long interval fire was characterized as a low- severity surface fire are indicated by an * symbol, and pairs where both fires were low- severity surface fire are indicated with a † symbol.

**Table 2.** ANOVA results for the effects of parameters of interest in individual models on bacterial community dissimilarity after controlling for soil horizon and drainage class

| Variable | Sum of Squares | F-value | p-value | R<sup>2</sup> |
| --- | --- | --- | --- | --- |
| Log difference in understory stems (all) | 0.007 | 5.79 | 0.023 * | 0.15 |
| Log difference in understory stems (conifer) | 0.056 | 4.47 | 0.043 * | 0.12 |
| Log difference in understory stems (broadleaf) | 0.044 | 3.45 | 0.073 | 0.14 |

### Short-Interval and Long-Interval differences in richness

Richness was greater in SI sites for some pairs (up to 93% greater richness in SI sites) and lower in SI sites for others (up to 53% lower richness in SI sites), but was not consistently significantly different (Figure 5, paired t-test, p = 0.81, p = 0.39, p = 0.66, and p = 0.96, for upland mineral soil, upland organic soil, wetland mineral soil, and wetland organic soil, respectively).

**Figure 5.**
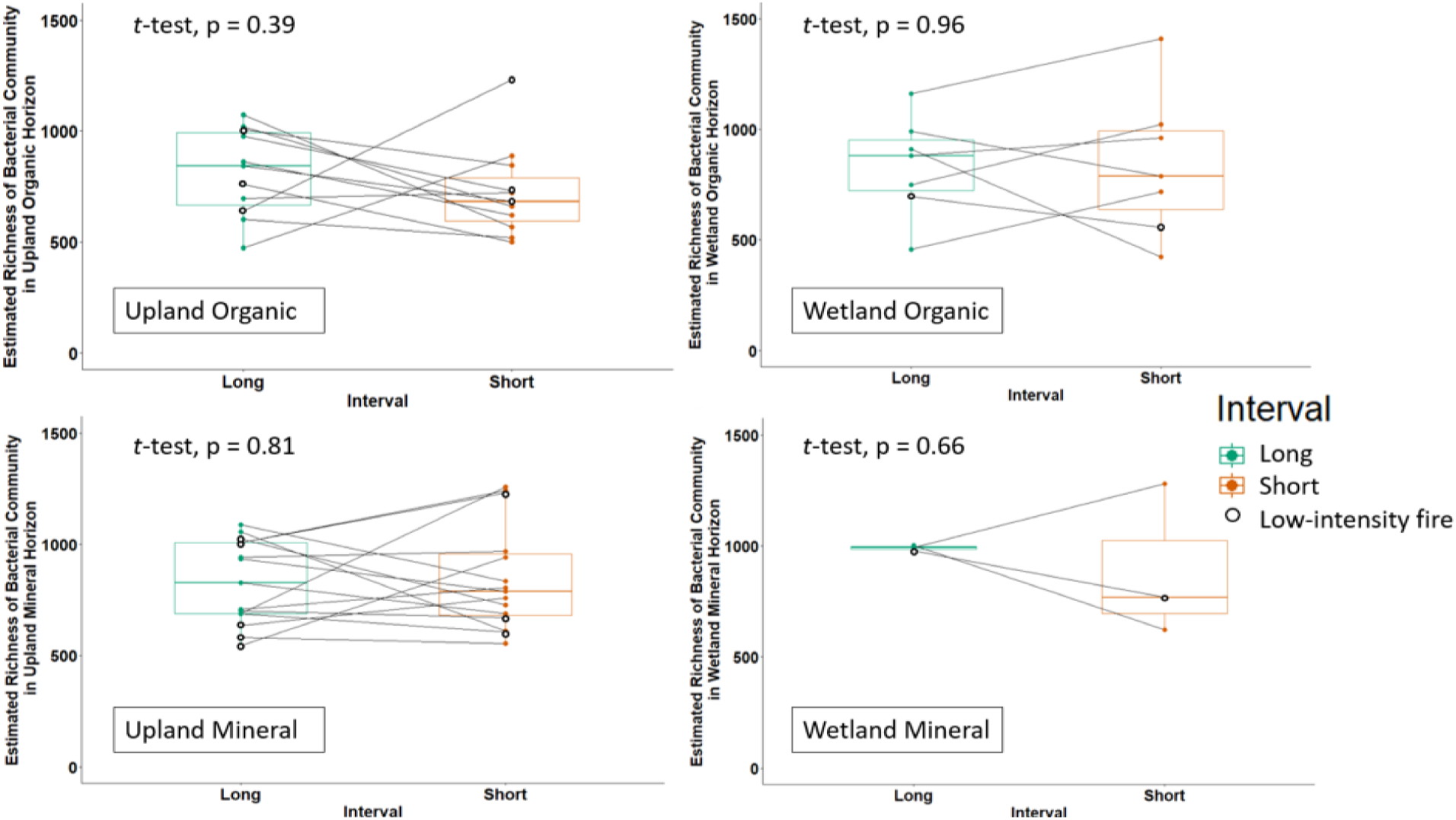
Boxplots of estimated bacterial community richness from paired (joined by grey lines) short and long fire free interval samples in the a) upland mineral soil, b) upland organic soil, c) wetland mineral soil, and d) wetland organic soil horizons (paired t-tests, p = 0.81, 0.39. 0.66, and 0.96, respectively). Each point represents a sample. Points outlined in black indicate a sample where fire was characterized as a low-severity surface fire.

None of our factors of interest had significant effects on bacterial community richness between paired SI and LI sites. Percent change in estimated richness between paired SI and LI samples was not associated with differences in FFI between paired sites (p = 0.08; Supplemental Table 5). Differences in estimated richness between paired SI and LI samples were also not associated with TSLF (p = 0.80, Supplemental Table 5), vegetation transitions (p = 0.33, Supplemental Table 5), difference in number of stems (p = 0.11, Supplemental Table 5), or drainage class (p = 0.93, Supplemental Table 5), after controlling for soil horizon and drainage class. Bacterial richness did not improve predictions of seedling recruitment, as estimated by live understory stem density (p = 0.32, Supplemental Table 6). After identifying differentially abundant bacteria (results below), we also wanted to know whether including differentially abundant taxa of interest would improve predictions of seedling recruitment. None of our top responding OTUs (identified as *Blastococcus, Rhizobiaceae,* and *Callaberonia*) significantly improved predictions of the recruitment of conifer and broadleaf seedlings in an ANOVA, after accounting for live overstory stems, site moisture, TSLF, interval, and OTU abundance as predictor variables.

### Differentially abundant bacteria between short interval and long intervals

We identified taxa that were differentially abundant in LI *vs*. SI sites in uplands (Figure 6a, Supplemental Figure 6) and wetlands (Figure 6b, Supplemental Figure 5). Of the most differentially abundant taxa, one taxon from the genus *Blastococcus* had 6.6 and 4.7 times greater relative abundance in SI sites than LI sites, in both uplands and wetlands, respectively, and was abundant across samples (uplands: 1.5% ± 1.6%; wetlands: 0.7% ± 1.2%). Of the taxa that were most differentially abundant in LI sites over SI sites, for uplands, we identified a taxon that had 3.9 times higher relative abundance in LI sites and was also relatively abundant overall (0.75% ± 0.51%) from the family *Burkholderiaceae*. For wetlands, we identified a taxon that was 5.0 times more abundant in LI sites and was relatively abundant overall (0.30% ± 0.26%) from the family *Rhizobiaceae*.

**Figure 6.**
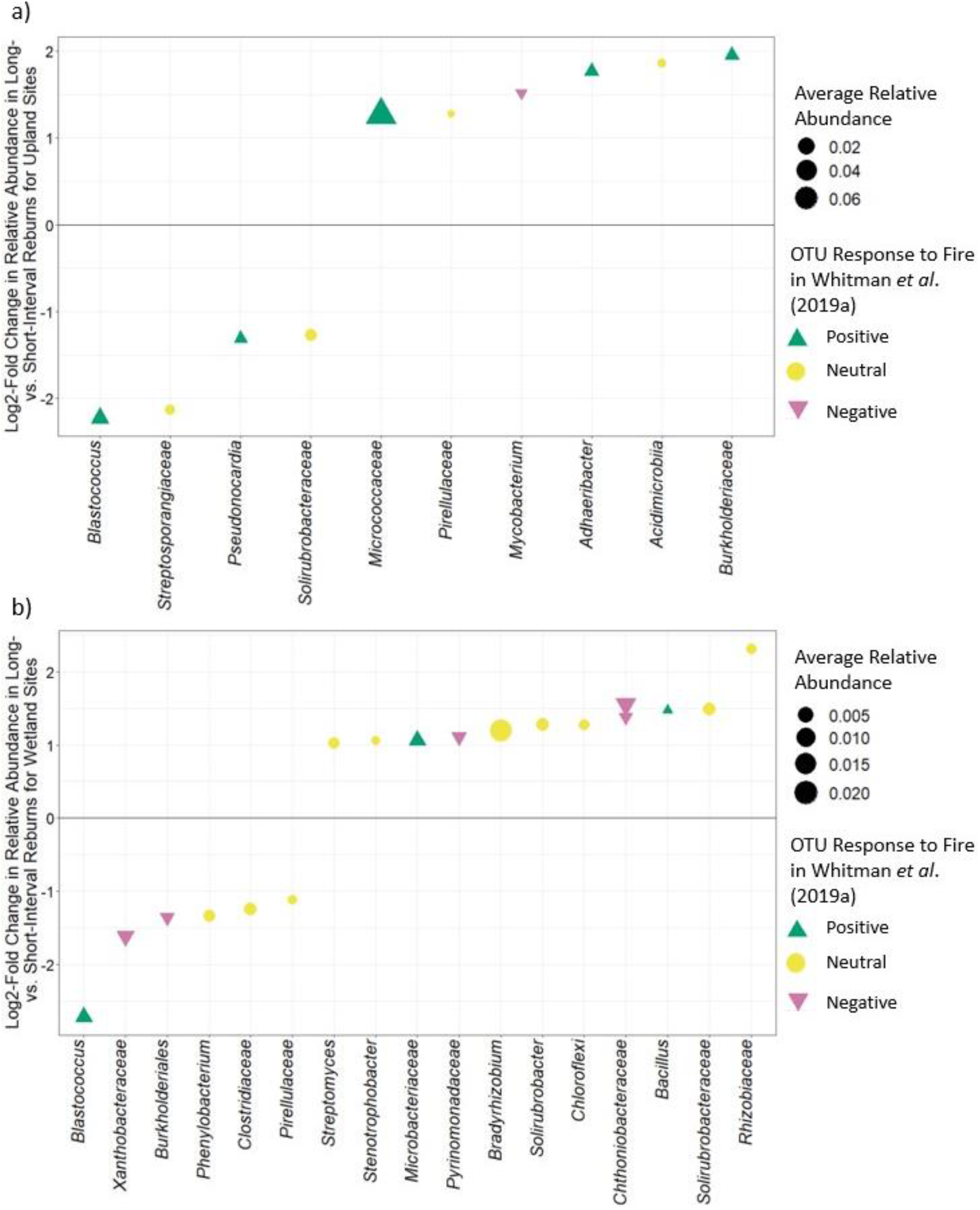
Log_2_-fold change in relative abundance in long *vs.* short fire-free intervals for (a) upland and (b) wetland sites, after controlling for Site ID and soil horizon. Each point represents a single OTU. Color and shape indicate the response the OTU had in Whitman *et al.* (2019a) (green triangle = positive response to fire yellow circle = neutral response to fire, pink upside-down triangle = negative response to fire,). The x-axis label indicates the finest-scale taxonomy available. Size of points is scaled by the average relative abundance of taxa in short- and long-interval sites. Only significantly differentially abundant taxa (p_FDR_<0.05) and responses greater than |1| are plotted. Solid line indicates no difference in relative abundance between long *vs*. short interval sites, therefore points above the line indicate taxa that were significantly more abundant in long-interval upland or wetland sites, and points below the line indicate taxa that were significantly more abundant in short-interval upland or wetland sites.

## Discussion

### Shorter fire-free intervals alter bacterial communities, reflecting changes in vegetation and soil properties

In addition to the clear differences in bacterial community composition between different drainage classes and soil horizons (Figure 2), short interval reburns also shifted bacterial community composition (Figure 3). The bacterial community changes between SI and LI were associated with decreased conifer dominance, as well as higher soil pH (Figure 3). This echoes the types of changes that have been seen in the aboveground communities in this region (Whitman *et al.*, 2019b), indicating that soil bacterial community composition will also be affected by predicted changes in this region – specifically, decreasing fire-free intervals. The extent to which changes in the bacterial communities are being driven by changes in the vegetation community, and vice-versa, is not possible to determine with this dataset and experimental design. As discussed in more detail below, the presence of vegetation transitions was not associated with greater differences in bacterial community composition between paired SI and LI sites, nor did we find a strong association between aboveground vegetation and soil bacterial communities. This lack of clear coupling may suggest that the primary drivers of changes to plant and bacterial communities with shorter fire-free intervals differ. While soil pH was not a significant predictor of community composition at the whole-dataset level after controlling for drainage class and soil horizon, higher pH was associated with shifts in community composition from LI to SI sites (Figures 2 and 3). This could be explained if short interval reburns were either higher severity, increasing total combustion and ash production, which would result in an increase in pH, particularly for combustion at higher temperatures (Bodí et al., 2014), or just simply from the compound effects of the two most recent fires, maintaining higher ash levels between the two. The shift toward increased soil pH in SI sites is notable, since soil pH is regularly found to be a powerful predictor of bacterial community composition in regional datasets (Rousk *et al*., 2010; Bahram *et al*., 2018). Given this observation, it would be interesting to determine whether pH shifts are the primary driver of shifts in bacterial community composition with shorter interval reburns. Since this was not our primary question for this study, future studies could be designed to test this hypothesis.

We found that the effects of short interval reburns on paired site dissimilarities were not clearly moderated by any of our predicted factors: TSLF, difference in FFI, vegetation transition, or the influence of drainage class. We had predicted that increasing TSLF would decrease Bray-Curtis dissimilarities between paired sites, as communities recover and converge on similar states post fire. If this had been the case, it might have suggested that bacterial communities are resilient to the effects of short interval reburns. However, our data do not indicate that increasing TSLF allows communities to converge meaningfully, over the range of TSLF studied (1-21 years). This is consistent with other studies of post-fire recovery in microbial communities, which have indicated that it can take decades to over a century for soil microbial communities to return to their pre-burn states (Dooley and Treseder, 2012; Ferrenberg *et al.,* 2013; Dove and Hart, 2017). Thus, it may not be surprising that the effects of short interval reburns did not seem to decrease community dissimilarity over the timescale of this study. That said, while we did not detect a significant effect, we might speculate that such an effect could still emerge over longer timescales. LI and SI pair dissimilarities at longer TSLF (12-21 years) ranged from 0.46-0.78 (more similar), while dissimilarities at shorter TSLF (1-2 years) ranged from 0.58 – 0.91 (less similar). While the ranges overlap, they do suggest a trajectory towards more similar communities after longer times since last fire. Future studies might be designed to target an even broader distribution of TSLF across the SI-LI pairs to directly test this question, or to trace these same sites over time. However, such a sample set may be difficult to collect, as it becomes increasingly difficult to conclusively identify FFI as we move beyond the range of modern satellite data records.

Since SI *vs*. LI was a significant predictor of bacterial community composition (Figures 2 and 3), we expected that communities might become increasingly dissimilar as differences in FFI increased. As differences in FFI increase, factors that could affect microbial community composition, such as soil properties or vegetation composition, may also become increasingly different, extending microbial community dissimilarities between the paired sites. However, larger differences in FFI were not associated with more dissimilar communities. This suggests that the differences in FFI do not scale consistently with differences in the direct or indirect effects of fire on microbial community composition. This may also point to the range of possible outcomes from a short interval fire. As a simplified example, there can be high-severity short interval fires where a recent burn has left high fuel loads, supporting significant loss of biomass and degradation of germination substrate, and leading to poor conifer generation. At the other extreme, there can also be low-severity short interval fires where frequent recent fires lead to surface fires with less tree mortality and SOM combustion, and safe germination sites. Thus, we might expect there to be no consistent association with increasing differences in FFI and bacterial community dissimilarity, as was observed here.

Vegetation can structure microbial communities through a wide range of mechanisms, such as interactions in the rhizosphere or through differences in litter inputs (Berg and Smalla, 2009; Merilä *et al*., 2010), so we had predicted that paired sites where differences between SI and LI sites were so great that leading tree species were different (“Vegetation Transition”) would also be associated with greater differences in bacterial community composition. Counter to our predictions, differences in the leading tree species between paired SI and LI sites were not associated with more dissimilar bacterial communities. Furthermore, differences in total vegetation cover or vegetation community dissimilarities were also not associated with bacterial community dissimilarity. These findings are somewhat consistent with our previous work in the region (Whitman *et al*., 2019a), where we found that understory vegetation and microbial community dissimilarities were not related for upland soils, which dominate this dataset. They are also consistent with the findings of Adkins et al. (2020) in a Sierra Nevada mixed conifer forest, who suggested that post-wildfire vegetation effects on community-level soil microbial properties were largely indirect and mediated through soil properties. However, when we considered a more sensitive measure of change in the understory vegetation community - difference in understory tree stem counts - this factor was significantly associated with increasing bacterial community dissimilarities (Table 2 and Figure 4). This metric allows us to discern sites with differences in tree regeneration, even if the leading species did not change. Differences in tree regeneration could have important effects on local soil conditions, including temperature and moisture, as sparsely vegetated sites or sites with more seedlings than mature trees may be warmer and drier (Hoecker *et al*., 2020). This finding raises the question, are bacterial communities different because there was more regeneration, or was tree regeneration more successful because of the bacterial community? Future experiments could be designed to investigate this question directly.

Although we expected that microbial communities in wetlands would be buffered from potential effects of short interval reburns more than uplands, we did not see an effect of drainage class on community dissimilarities between paired LI and SI sites. Because sites were sampled more than one year post fire, we were not able to generate accurate estimates of fire severity, which is known to affect bacterial communities (Whitman *et al*., 2019a; Day *et al*., 2019). Our simplified metric for inferring low-severity surface fires does not apply well here either, since trees are generally less abundant in the wetland sites. However, if some wetlands had high burn severities for short interval reburns, then we would not have expected our hypothesis to be supported. To examine this possibility, we can consider two contrasting wetland pairs, which were both sampled 8 years post fire. The first pair has the greatest dissimilarity between SI and LI communities (Bray-Curtis dissimilarity 0.901 in the mineral soil and 0.827 in the organic soil). For this pair, the organic horizon depth in SI was 88.45% lower than LI, and the dominant tree type transitioned from white spruce to balsam poplar. These metrics may suggest a higher severity fire in the SI. In contrast, the organic and mineral soil communities of the second pair were more similar (Bray-Curtis dissimilarities of 0.699 and 0.575, in mineral and organic horizons, respectively); in this pair, the organic depth in SI was only 19.45% lower than LI, and the leading tree type did not change (trembling aspen-mixed forest). These metrics may suggest a lower severity fire in SI compared to the first pair. The possibility that the first pair experienced a relatively high severity fire in the SI (indicated by a greater difference in O horizon thickness and change in leading tree type) may explain why the paired samples are more dissimilar, and would be counter to our hypothesis that the wetlands would experience lower severity fires across the board, leading to smaller effects of short interval reburns. Relative to severely burned SI uplands, the biogeophysical differences between SI and LI wetlands may be lower (*e.g.*, thicker residual organic horizons and greater post-fire conifer dominance; Whitman *et al.* 2019b), but repeated wildfires nonetheless produce meaningful differences in both above and belowground communities.

### Bacterial richness is not consistently affected by shorter fire free intervals

We expected that bacterial richness would not be clearly affected by short interval reburns, largely because soil bacterial communities are extremely rich to begin with and, despite advances in sequencing technologies, methods for accurate richness estimation in high-throughput amplicon sequencing datasets remain limited (Willis, 2019). This expectation was supported - although most LI sites were richer than their SI pairs (Figure 5), this difference was not significant and the percent difference in richness ranged widely - from 93% higher (SI richer than LI) to 53% lower (LI richer than SI). Although fires often reach temperatures at the soil surface that would be expected to kill all bacteria (Pingree and Kobziar, 2019), heating attenuates rapidly with depth in the soil, meaning that lethal temperatures in the surface of the soil may be accompanied by almost no changes just a few centimeters below the surface. Thus, it may not be surprising that the effects of fire on bacterial richness have been inconsistent across studies, with some studies reporting a decrease in bacterial richness with increasing fire severity (Sáenz de Miera *et al.*, 2020), or reporting a lack of significant changes (Pressler *et al.* 2019; Whitman *et al*., 2019a). Our findings indicate that, although SI *vs*. LI have different bacterial community compositions (Figures 2 and 3), this difference is likely driven more by changes in the overall community structure or by a few community members, rather than significant differences in richness. While richness has often been linked to ecosystem multifunctionality (Delgado-Baquerizo *et al.*, 2017) or resistance and resilience (Shade *et al.*, 2012), these relationships are not always straightforward in soil communities, which generally have high functional redundancy (Nunan et al., 2018). This functional redundancy is perhaps also reflected in our finding that including bacterial richness in our models did not improve predictions of understory stem regeneration. Thus, even for the paired sites where bacterial richness was lower in SI than LI, we should not necessarily predict functional limitations.

### Short- and long-interval communities have distinct responding bacteria

While the fire ecology of plants, and even many fungi, is well-characterized and has long been studied (Cooper, 1961; Seaver, 1909), the bacterial equivalents of fire responder species like *Pinus banksiana* or the fungus *Pyronema* are only just being established. By identifying specific bacteria associated with SI or LI sites, we sought to both expand our understanding of fundamental questions about fire ecology for bacteria, as well as to offer potential hypotheses about the effects that changes to soil bacterial communities might have on ecosystem functioning. For example, taxa that are more abundant in SI sites may be able to thrive under the conditions characteristic of frequently burned areas. Thus, we might predict these microbes would exhibit traits that allow them to consume pyrogenic organic matter more effectively, thrive in areas that are more drought-prone, or be common in areas with less vegetation cover and tree regeneration. Taxa that are more abundant in LI sites could be general fire responders that are characteristic of normal fire regimes; these taxa may thrive in post-fire environments, just not when their habitat is significantly altered by short interval reburns.

Even though our study sites spanned a wide range of post fire conditions, we were able to identify several taxa that were consistently differentially abundant in SI and LI sites. Because of this experimental design, we believe that these observed responses may be robust within this region, rather than site-specific anomalies. Our most prominent responder was a *Blastococcus* OTU, which was the most abundant responding OTU in both wetlands and in uplands (Figure 6). It was over four times more abundant in SI soils than in LI soils, and consistently represented a large proportion of the total community across sites. This was the same OTU (100% identical over the sequenced region) as an OTU in Whitman *et al*. (2019a) that was enriched in burned sites and was increasingly abundant in sites with increasing fire severity. In holm-oak forests of Spain, members of *Blastococcus* also increased in abundance after wildfires in rhizosphere soils *versus* neighboring unburned areas (Fernández-González *et al.*,2017; Cobo-Diaz *et al.*, 2015). The sequence for this taxon was also 100% identical to a recently classified *Blastococcus deserti* sp. isolated from a desert sample (Yang *et al.,* 2019). While clearly a fundamentally different ecosystem, in their study, Yang *et. al.* noted that this isolate was able to survive at temperatures up to 50 °C and utilize D-salacin as a C source, which is a compound found in the bark of *Populus* and *Salix* species (Palo, 1984), when other related *Blastococcus* strains in their study could not. Both of these traits offer mechanisms that could help this taxon thrive in the post-fire environment: its potential ability to survive high temperatures during fires, and also metabolize compounds characteristic of the organic horizon of an aspen-dominated forest – a common successional species after short interval reburns in this region (Whitman *et al*., 2019b; Johnstone and Chapin, 2006). Together, these studies highlight traits that allow us to propose this *Blastococcus* OTU as a classical “pyrophilous” bacterium, which thrives in burned soils and even becomes increasingly abundant with more frequent and more severe burns.

For wetland sites, the most prominent LI associated taxon was classified as belonging to the family *Rhizobiaceae*. This OTU was detected in the Whitman *et al*. (2019a) study, but did not show a negative or positive response to fire in that dataset. Members of family *Rhizobiaceae* are often associated with nitrogen fixation (Spaink, Kondorosi, and Hooykaas, 1998), which could be an important source of N in post-fire ecosystems (Smithwick et al., 2005). Some members of the *Rhizobiaceae* family also perform other steps of the N cycle, including denitrification (Rich *et al*., 2003). While these roles can be seen to be relevant post-fire, it is not immediately clear why these taxa were more enriched in the LI wetland sites. One possibility is that if the SI wetland sites were drier, then the LI sites may have had conditions more conducive to denitrification (low oxygen due to high moisture, and high organic matter (Martínez-Espinosa *et al*., 2021)) that selected for these taxa. Similarly, N fixation could also be more likely to be supported under these conditions: it is a highly energy intensive process requiring both lots of C for energy and at least locally anoxic conditions to protect the oxygen-sensitive nitrogenase enzyme (Smercina *et al*., 2019). However, even though this *Rhizobiaceae* OTU is 100% identical over the sequenced region to taxa that are known to be able to perform these functions, there may still be important functional differences. We should consider these speculations on their role in post-fire N cycling at LI sites as hypotheses for future testing.

For upland sites, the most prominent LI associated taxon, identified as being from the *Burkholderiaceae* family, also matched an OTU in Whitman *et al.* (2019a) that was found to be more abundant in burned sites, to the point of not being detected at any unburned sites. Upon first consideration, this seems somewhat surprising – why would a positive burn responder be more abundant at LI burned sites, whose soil properties would presumably be less influenced by fire? In considering this question, it is important to remember that all of the sites in this study were burned, so we should not necessarily expect that LI-associated taxa should also be classified as negative burn responders. Furthermore, the dataset in Whitman *et al*. (2019a) spanned a wide range of burned sites, including both long and short interval reburns across a range of severity and vegetation communities. One possible explanation for this association could be that members of the genus *Burkholderia*, from this same family, have been found to be more abundant in lower pH soils (Stopnisek *et al*., 2014), and we observed here that short interval reburns were associated with higher pH (Figure 3). Supporting this observation, the corresponding OTU from Whitman *et al*. (2019a) clustered in a module associated with fire-responsive taxa that were more abundant at lower pH sites in a co-occurrence network.

This LI-associated organism may also play an important role in post-fire plant establishment. While the finest-scale matching taxonomy in the SILVA database placed this LI-associated OTU within the *Burkholderiaceae* family, using NCBI BLAST, we found it is a 100% ID match for a *Caballeronia sordidicola* that was isolated from a lichen in Svalbard Archipelago (AF512827.1; Kim *et al*., 2017) and for a *Caballeronia sordidicola* spruce tree endophyte from a sub-boreal forest in British Columbia, Canada (MG561776.1; Puri *et al*., 2018). Furthermore, the authors of this second study identified that this isolate has the capacity to fix nitrogen (Puri *et al*., 2018), readily colonized pine and spruce seedlings, increasing their biomass production 4-7-fold (Puri *et al*., 2020a) and provided more than 50% of both lodgepole pine (*Pinus contorta*) and white spruce seedlings’ N requirements (Puri *et al*., 2020b). This is strongly consistent with our observation that conifer seedlings were more abundant at LI sites. While our study design does not allow us to separate the cause and effect between the increased presence of putative endosymbiont diazotroph *Callaberonia sordidicola* and more abundant conifer seedlings at LI sites, it clearly raises pressing questions. For example, linking our observations, we might ask whether increases in pH due to short interval reburns shift the soil environment away from conditions that are optimal for *Callaberonia sordidicola*, exacerbating other factors that contribute to poor conifer seedling establishment, such as depleted seed banks or suboptimal seedbed conditions associated with short interval reburns. Certainly, designing experiments to probe the ecological implications of the depletion of this potentially highly influential spruce and pine endosymbiont in short interval reburns would be an important next step for this system.

## Conclusions

Short interval reburns result in altered soil bacterial communities. These altered short-interval communities were associated with sites that had fewer understory stems, a lower percentage of conifer seedlings, and higher pH. These findings suggest that, as has been seen in plant communities, bacterial communities are also likely to change as fire-free intervals decrease. Although there were changes in overall community structure, we did not see consistent effects of short interval reburns on bacterial richness. We did, however, identify specific taxa that were characteristic of SI *vs.* LI sites, including the abundant and strongly SI-associated *Blastococcus* OTU. This common successional taxon in uplands as well as wetlands after short interval reburns is a general fire-responder in this region and in other parts of the world. Beginning to parse its specific fire ecology, future experiments could determine whether this is due to an ability to survive higher temperatures and an ability to metabolize compounds characteristic of the organic horizon of an aspen-dominated forest. Additionally, the depletion of diazotrophic conifer endophyte *Callaberonia sordidicola* in short interval reburn sites raises questions about whether its depletion is contributing to – or merely reflects – poor conifer seedling recolonization post-fire at short-interval reburns.

There are numerous other future directions for this work. First, directly investigating some of the hypotheses generated here is of interest, including the extent to which bacterial community shifts due to shorter fire-free intervals are driven by changes in soil pH. Research is needed to elucidate the relative influence of post-fire microbial communities in determining vegetation community development and facilitating or impeding tree recruitment and, conversely, further research is needed to identify the influence of post-fire vegetation on soil microbial community development and the strength of these interacting effects. Additionally, while identifying specific fire-responsive taxa is a critical first step in developing a fire ecology framework for bacteria, the next steps will be to investigate the functional traits of these responding taxa in a mechanistic way, in order to further understand their individual ecology and what effects changes in their populations might have on ecosystem processes.

## Acknowledgements

The field campaign in which these data were gathered was funded by the government of the Northwest Territories, and the Natural Sciences and Engineering Research Council of Canada (Funding Reference Number: CGSD3-471480-2015). Environment and Climate Change Canada, Sam Haché, Parks Canada Agency, and Jean Morin provided in-kind support. Matthew Coyle, Kathleen Groenwegen, Xianli Wang, Mary Stephens, Scott L. Stephens, Rodrigo Campos-Ruiz, and Josh Gauthier provided indispensable help in the field. This research was performed using the computing resources and assistance of the UW-Madison Center for High Throughput Computing (CHTC) in the Department of Computer Sciences. The CHTC is supported by UW-Madison, the Advanced Computing Initiative, the Wisconsin Alumni Research Foundation, the Wisconsin Institutes for Discovery, and the National Science Foundation, and is an active member of the Open Science Grid, which is supported by the National Science Foundation and the US Department of Energy’s Office of Science. T.W. and J. W. were partially supported by the U.S. Department of Energy (DE-SC0020351). The authors have no conflicts of interest to declare.

## Supplemental Information

Supplemental information associated with this study is available. Code used for all analyses can be found at https://github.com/WhitmanLab/WoodBuffalo2016. All sequences are deposited in the NCBI SRA under accession numbers [to be submitted upon paper acceptance].

