## Supplemental Information for "Short-interval reburns in the boreal forest alter soil bacterial communities, reflecting increased pH and poor conifer seedling establishment"

TITLE

Supplemental Table 1. PERMANOVA for parameters of interest on the Bray-Curtis dissimilarity of the paired sites

| Variable | Sum of Squares | F-value | p-value | $R^2$ |
|---|---|---|---|---|
| Pair ID | 10.927 | 1.89 | 0.001 * | 0.43 |
| Interval | 0.759 | 2.67 | 0.002 * | 0.03 |

Supplemental Table 2. PERMANOVA results for parameters of interest on the Bray-Curtis dissimilarity of the paired sites

| Variable | Sum of Squares | F-value | p-value | $R^2$ |
|---|---|---|---|---|
| Drainage Class | 1.606 | 5.54 | 0.001 * | 0.06 |
| Soil Horizon | 1.523 | 5.25 | 0.001 * | 0.06 |
| pH | 21.174 | 1.11 | 0.28 | 0.84 |

Supplemental Table 3. ANOVA results for parameters of interest in individual models for their effects on paired site Bray-Curtis dissimilarities after controlling for drainage class and soil horizon. Negative adjusted $R^2$ should be interpreted as 0.

| Variable | Sum of Squares | F-value | p-value | $R^2_{adj}$ |
|---|---|---|---|---|
| TSLF at time of sample | 0.025 | 1.839 | 0.18 | 0.04 |
| Vegetation Transition | 0.002 | 0.14 | 0.71 | -0.01 |
| Drainage Class | 0.032 | 2.37 | 0.13 | 0.04 |
| Difference in FFI | 0.035 | 2.626 | 0.12 | 0.07 |

Supplemental Table 4. ANOVA results for parameters of interest in individual models for their effects on paired site Bray-Curtis dissimilarities after controlling for drainage class and soil horizon.

| **Variable** | **Sum of Squares** | **F-value** | **p-value** | $R^2_{adj}$ |
|---|---|---|---|---|
| Difference in stem count | 0.074 | 6.256 | 0.02* | 0.16 |
| Difference in veg cover | 0.000 | 0.14 | 0.91 | -0.01 |
| Vegetation dissimilarity | 0.000 | 0.003 | 0.96 | -0.01 |

Supplemental Table 5. ANOVA results for parameters of interest in individual models for their effects on the percent change in richness of bacteria between paired sites in individual models after controlling for drainage class and soil horizon.

| **Variable** | **Sum of Squares** | **F-Value** | **p-value** | $R^2_{adj}$ |
|---|---|---|---|---|
| Difference in FFI | 4987 | 3.330 | 0.08 | 0.00 |
| TSLF at time of sample | 112 | 0.067 | 0.80 | -0.11 |
| Vegetation transition | 1563 | 0.972 | 0.33 | -0.07 |
| Difference in total stem count | 4256 | 2.797 | 0.10 | -0.02 |
| Drainage Class | 11 | 0.007 | 0.93 | 0.06 |

Supplemental Table 6. ANOVA results for parameters of interest on the total live understory stem count

| **Variable** | **Sum of Squares** | **F-Value** | **p-value** |
|---|---|---|---|
| Live overstory stems | 13340867 | 1.131 | 0.29 |
| Site moisture | 41102872 | 3.485 | 0.07 |
| TSLF | 97760904 | 8.288 | 0.005 * |
| FFI | 7062262 | 0.599 | 0.44 |
| Bacterial richness | 11983669 | 1.016 | 0.32 |

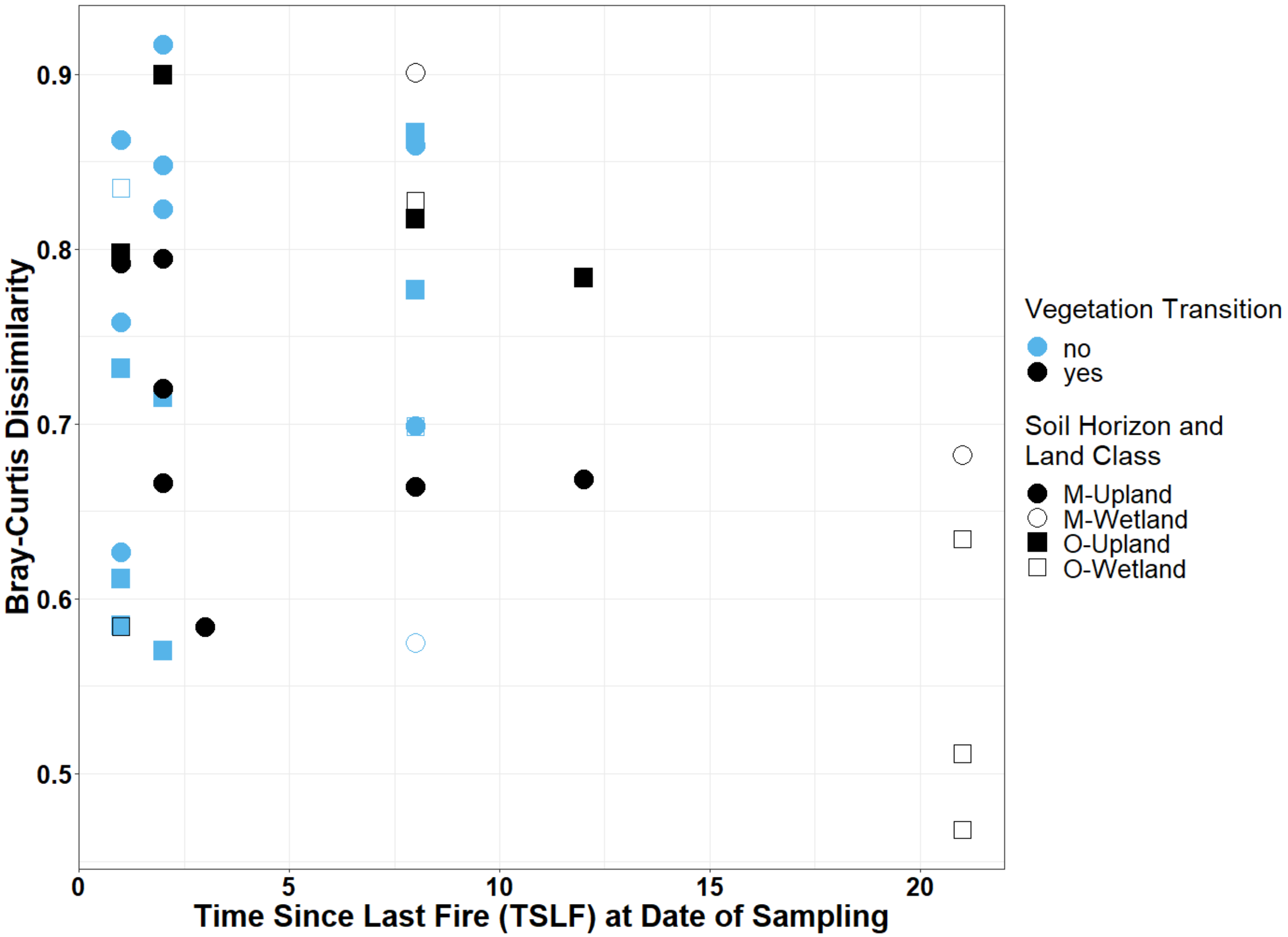

Supplemental Figure 1. Bray-Curtis dissimilarity of the bacterial community compositions between paired short and long-interval sites *vs.* time since last fire at sampling time. Vegetation transition (different leading species between paired plots) is indicated by color (light = no, dark = yes), land class is identified by fill (wetland = open symbols, upland = filled symbols), and soil horizon is identified by shape (mineral (M) = circle, organic (O) = square).

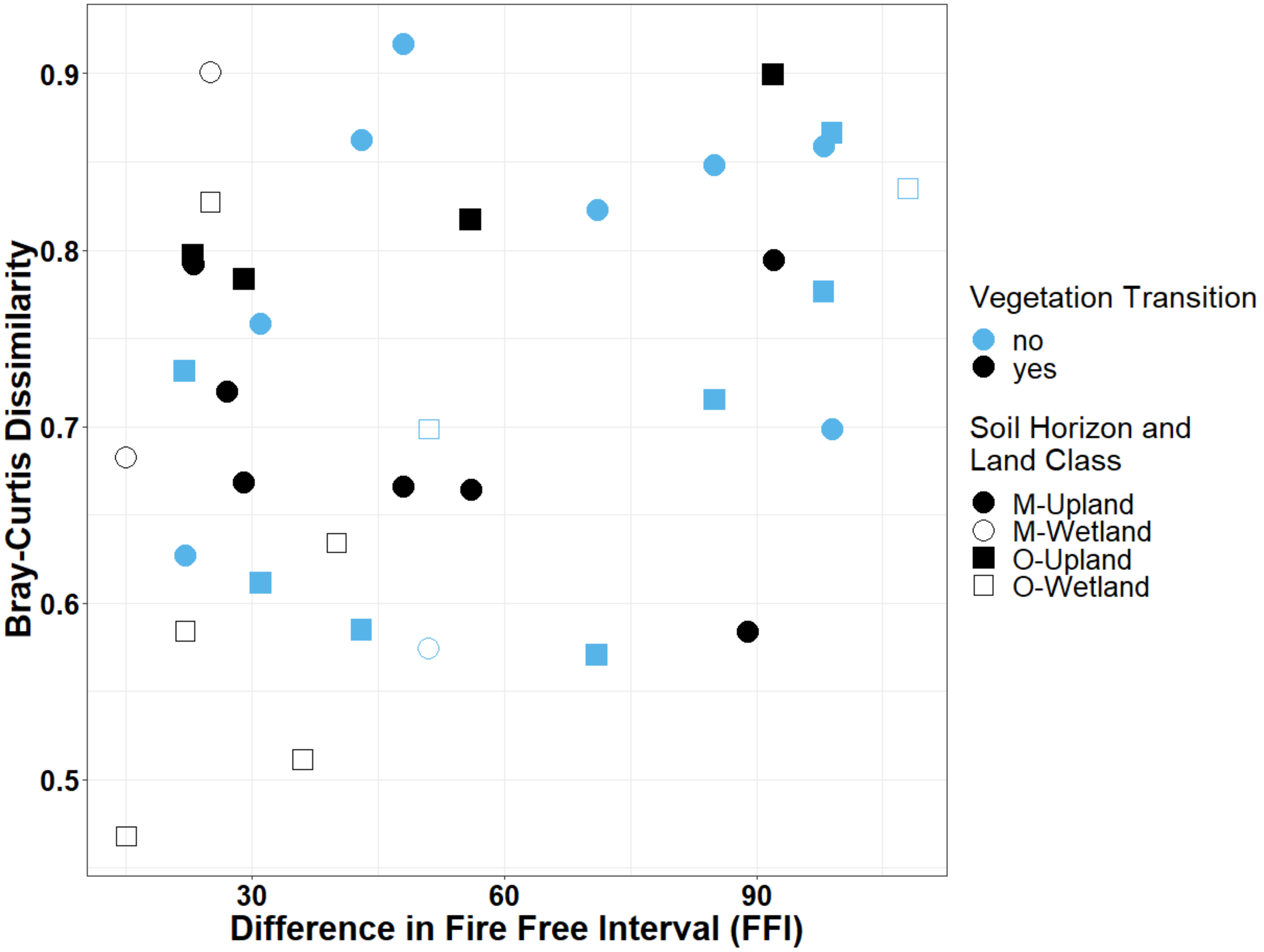

Supplemental Figure 2. Bray-Curtis dissimilarity of the bacterial community compositions between paired short- and long-interval sites *vs.* difference in fire free interval. Vegetation transition (different leading species between paired plots) is indicated by color (light = no, dark = yes), land class is identified by fill (wetland = open symbols, upland = filled symbols), and soil horizon is identified by shape (mineral (M) = circle, organic (O) = square).

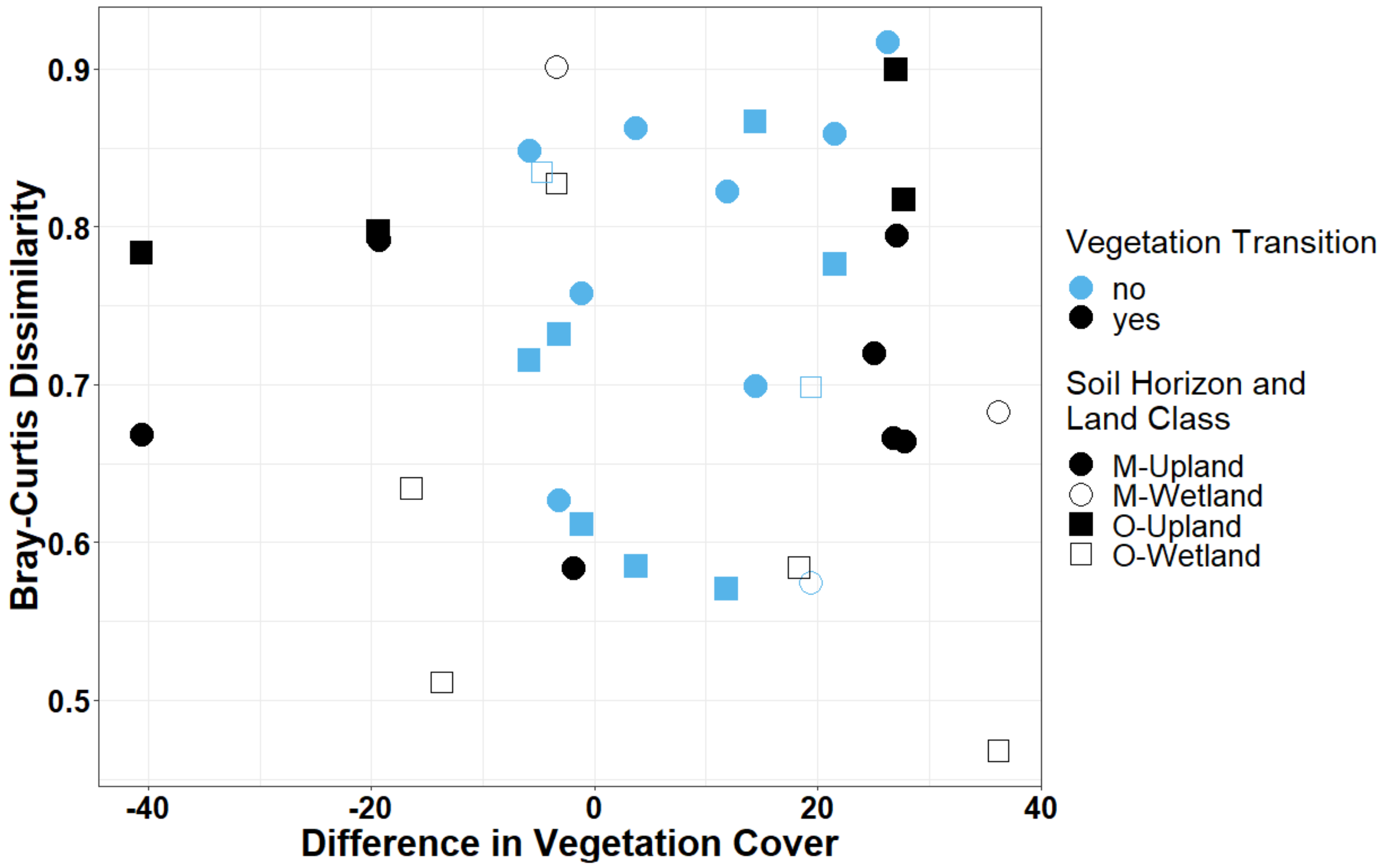

Supplemental Figure 3. Bray-Curtis dissimilarity of the bacterial community compositions between paired short and long-interval sites *vs.* the difference in vegetation cover in long interval – short interval sites. Vegetation transition (different leading species between paired plots) is indicated by color (light = no, dark = yes), land class is identified by fill (wetland = open symbols, upland = filled symbols), and soil horizon is identified by shape (mineral (M) = circle, organic (O) = square).

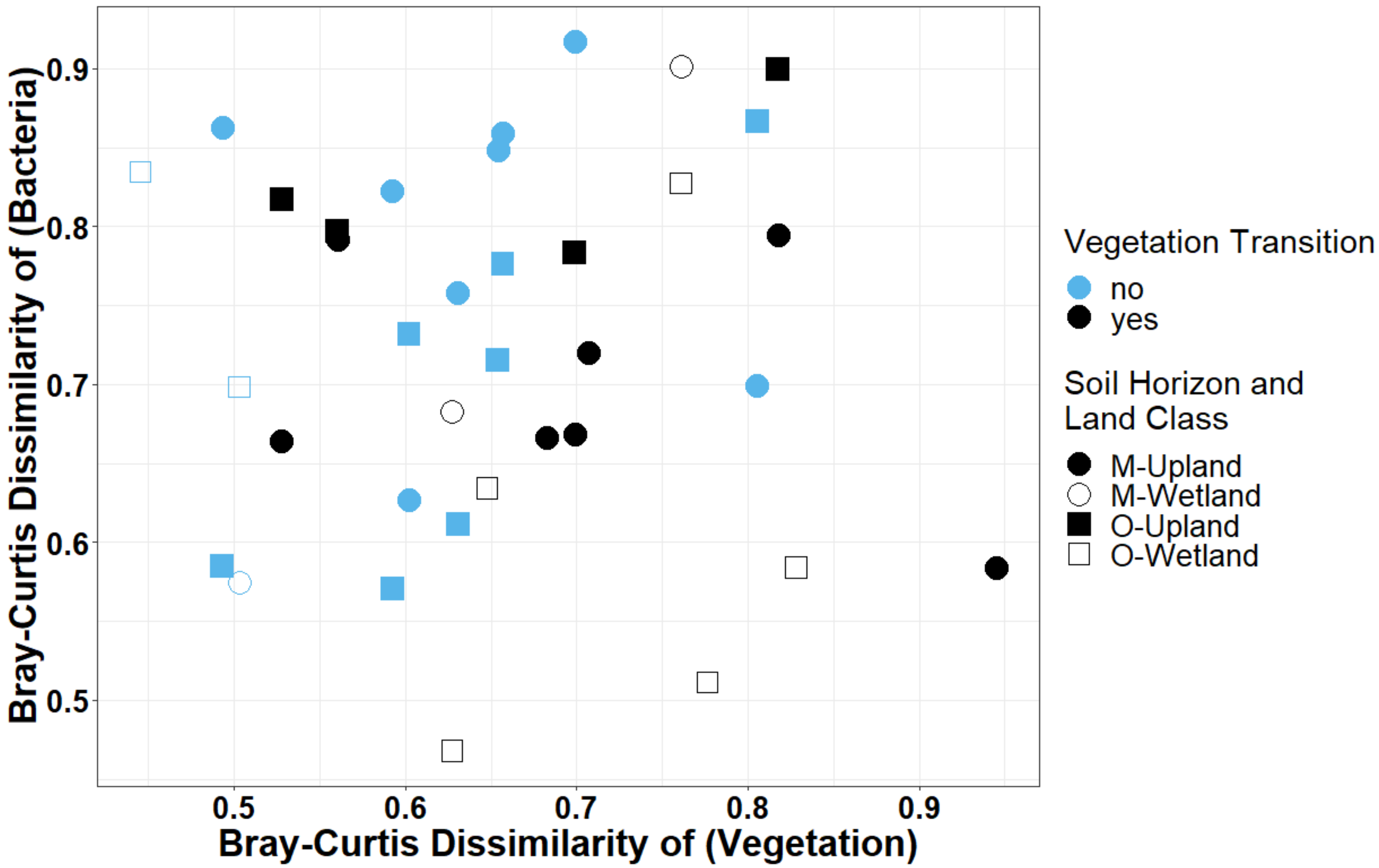

Supplemental Figure 4. Bray-Curtis dissimilarity of the bacterial community compositions between paired short and long-interval sites *vs.* Bray-Curtis dissimilarity of the vegetation community compositions between paired short and long-interval sites. Vegetation transition (different leading species between paired plots) is indicated by color (light = no, dark = yes), land class is identified by fill (wetland = open symbols, upland = filled symbols), and soil horizon is identified by shape (mineral (M) = circle, organic (O) = square).

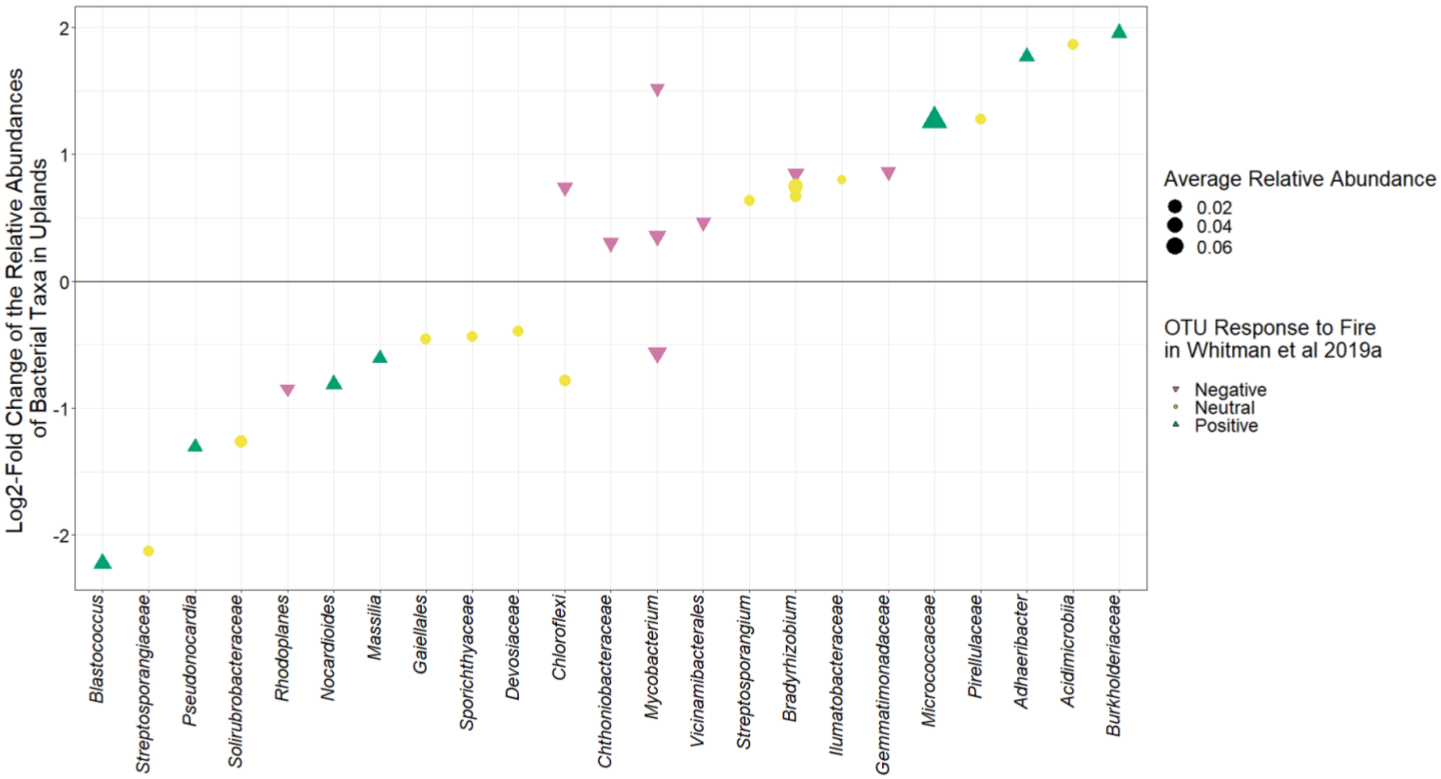

Supplemental Figure 5. $Log_2$-fold change in relative abundance in long-interval *vs.* short-interval reburns for upland sites, after controlling for Site ID and soil horizon. Each point represents a single OTU. Color and shape indicates the response the OTU had in Whitman *et al.* 2019a (pink upside-down triangle = negative response to fire, yellow circle = neutral response to fire, green triangle = positive response to fire). The x-axis label indicates the finest-scale taxonomy available. Size of points is scaled by the average relative abundance of taxa in short- and long-interval sites. Only significantly differentially abundant taxa ($p_{FDR}<0.05$) are plotted. Solid line indicates no difference in relative abundance between long vs. short interval sites, so points above the line indicate taxa that were significantly more abundant in long-interval upland sites, and points below the line indicate taxa that were significantly more abundant in short-interval upland sites.

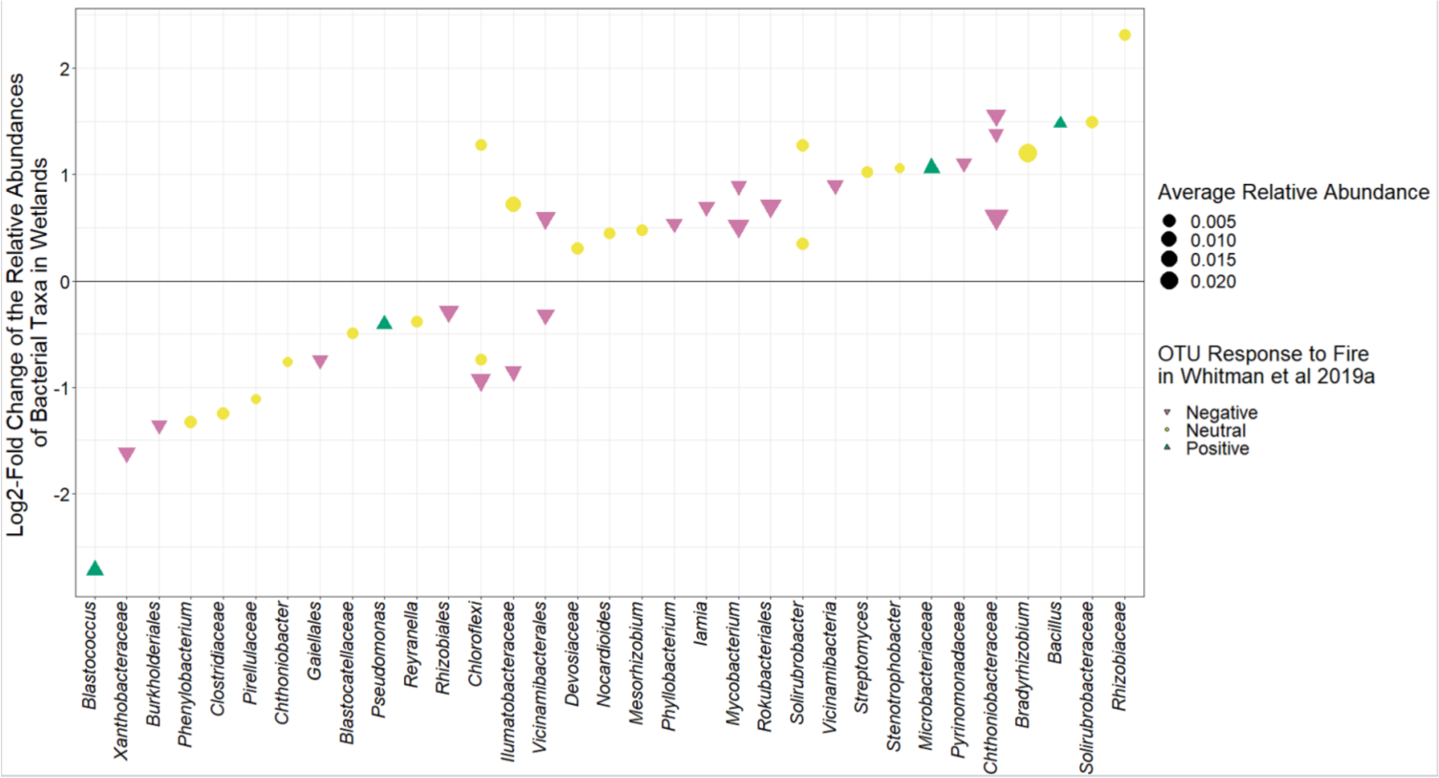

Supplemental Figure 6. $Log_2$-fold change in relative abundance in long-interval *vs.* short-interval reburns for wetland sites, after controlling for Site ID and soil horizon. Each point represents a single OTU. Color and shape indicates the response the OTU had in Whitman *et al.* 2019a (pink upside-down triangle = negative response to fire, yellow circle = neutral response to fire, green triangle = positive response to fire). The x-axis label indicates the finest-scale taxonomy available. Size of points is scaled by the average relative abundance of taxa in short- and long-interval sites. Only significantly differentially abundant taxa ($p_{FDR}$<0.05) are plotted. Solid line indicates no difference in relative abundance between long vs. short interval sites, so points above the line indicate taxa that were significantly more abundant in long-interval wetland sites, and points below the line indicate taxa that were significantly more abundant in short-interval wetland sites.

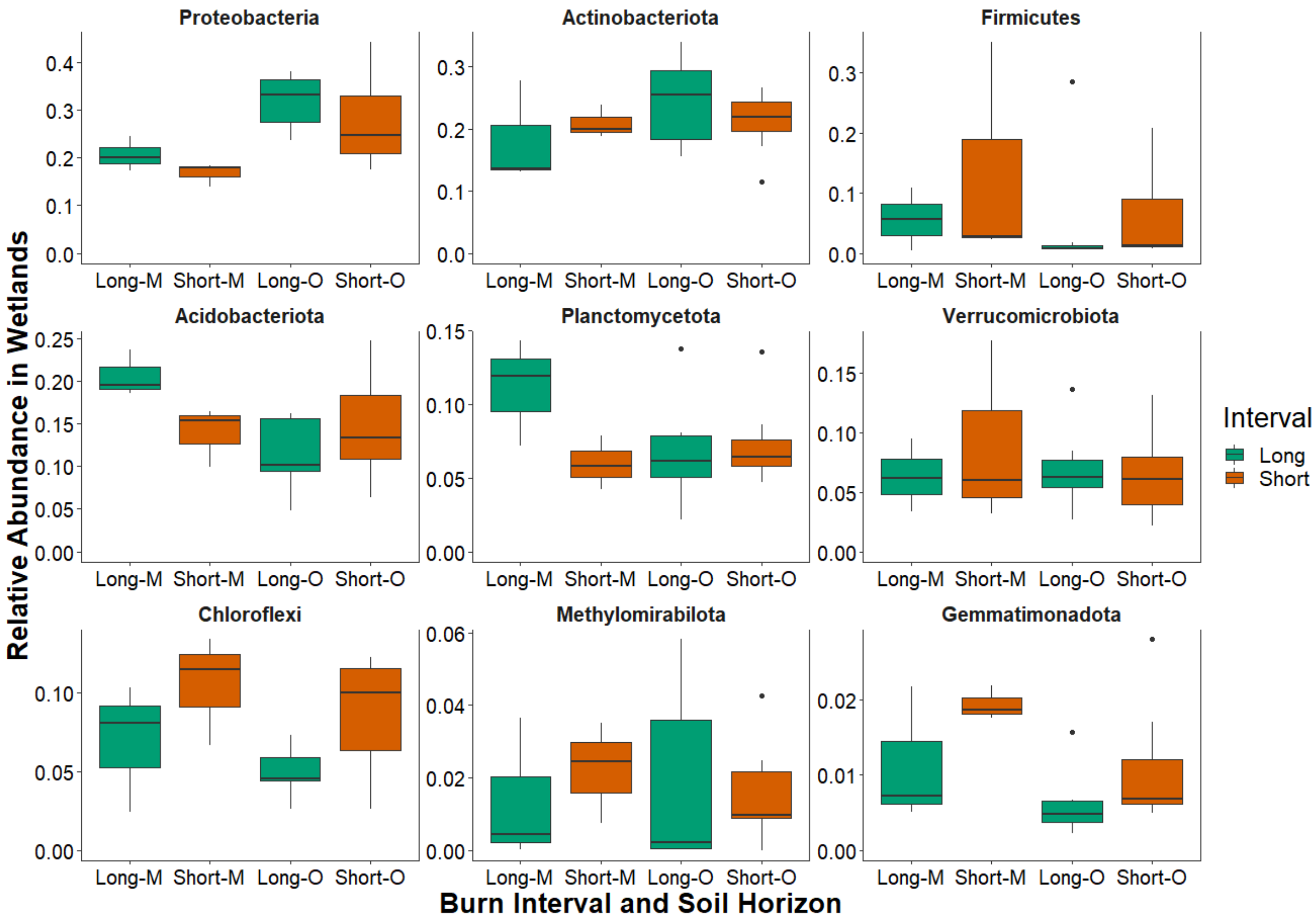

Supplemental Figure 7. Relative abundances of the top nine most abundant phyla in wetland sites. Taxa are grouped by burn interval (long = green, short = orange) and soil horizon (mineral = M, organic = O).

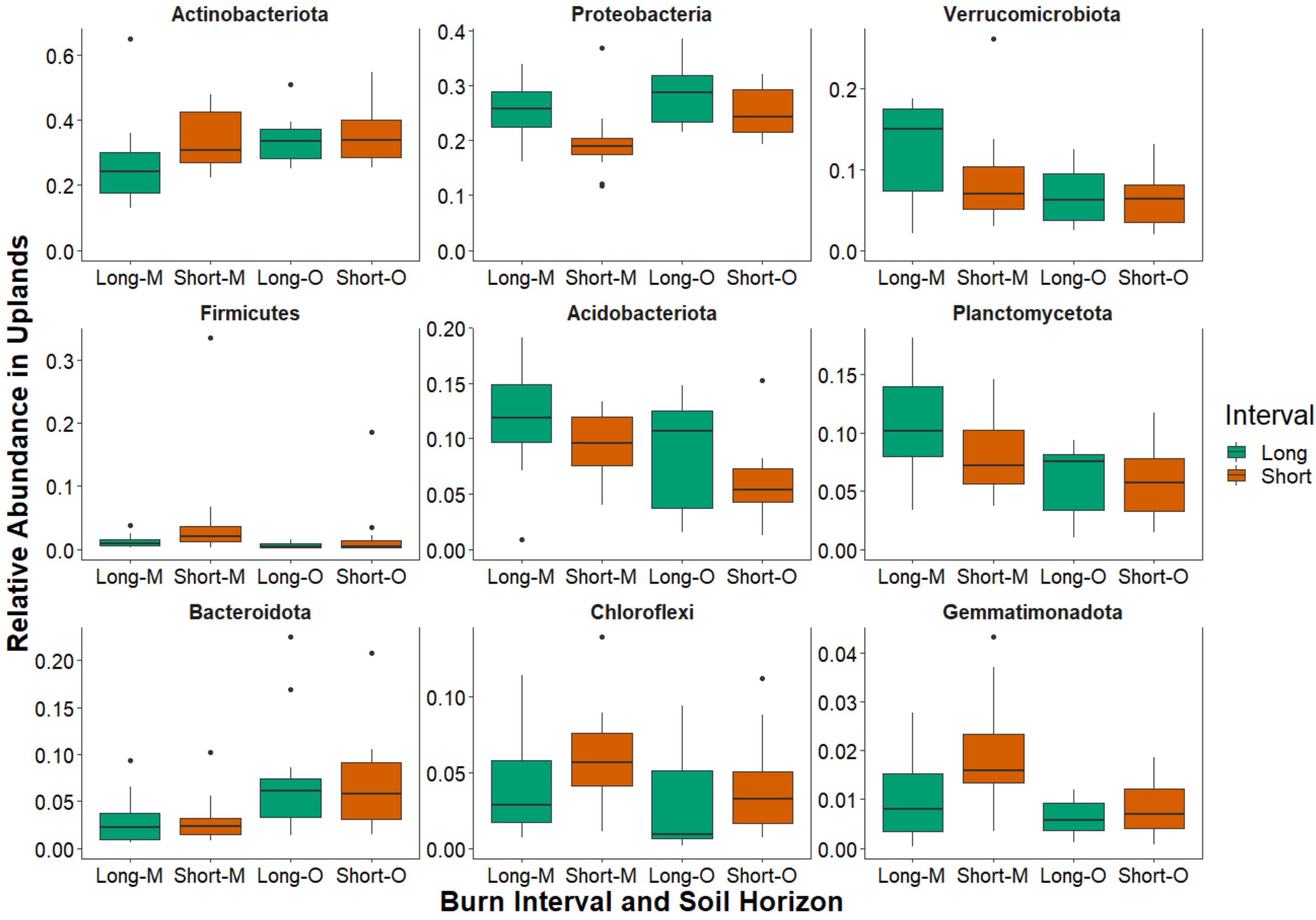

Supplemental Figure 8. Relative abundances of the top nine most abundant phyla in upland sites. Taxa are grouped by burn interval (long = green, short = orange) and soil horizon (mineral = M, organic = O).
